# Structural dynamics of sphingosine kinase 1 regulation and inhibition

**DOI:** 10.1101/2025.10.09.681406

**Authors:** Baharak Abd Emami, Ahmed Shubbar, Hope Woods, Mahmoud Moradi, Reza Dastvan

## Abstract

Sphingosine kinase 1 (SK1) generates sphingosine-1-phosphate, a bioactive lipid implicated in cancer and other diseases. Despite its clinical importance, the structural and dynamic basis of SK1 regulation and inhibition remains poorly understood. Using integrated spectroscopic and computational approaches, we uncover conformational transitions that govern substrate entry, catalysis, and inhibitor binding. Phosphorylation of Ser225 reconfigures the regulatory loop and reshuffles salt bridges, priming SK1 for membrane engagement and catalytic activity. We identify a previously uncharacterized catalytic intermediate with a distinct conformation and a highly dynamic lipid-binding loop 1 (LBL-1), sensitive to potent inhibitors such as PF-543. These inhibitors not only stabilize non-catalytic states but also induce LBL-mediated dimerization, blocking membrane binding and substrate access. Our findings reveal a multilayered regulatory mechanism driven by structural flexibility and establish a novel inhibitory paradigm. This framework provides critical insight into SK1 regulation and a foundation for developing next-generation SK1-targeted therapeutics.

## Introduction

The bioactive lipid sphingosine-1-phosphate (S1P) plays a key role in regulating mammalian cell growth, survival, and migration^1^. It is essential for lymphocyte trafficking, immune responses, vascular and embryonic development, and bone homeostasis^2–4^. S1P is produced intracellularly by sphingosine kinase (SK) isoform 1 and 2, then released extracellularly to carry out its (patho)physiological functions^5,6^. Aberrant accumulation of S1P is linked to cancer progression and other diseases, including atherosclerosis, diabetes, and inflammatory disorders^2,4,5,7–12^. ERK1/2 and PKC activation under hypoxic conditions, along with SK1 phosphorylation, induces conformational changes that facilitate membrane recruitment^13,14^. While a few SK1 crystal structures were solved over a decade ago^15–17^, key regulatory regions like the C-terminal tail and in some cases the loop bearing the phosphorylation site, Ser225, are missing or dissociated from the core protein. These regions are proposed to regulate conformational dynamics and the membrane recruitment^4,14^. Although existing structures provide valuable insights into the catalytic function and regulation of this important enzyme^4^, experimental validation using the full-length protein is needed to connect structural insights with functional dynamics. Thus, a comprehensive understanding of SK1’s structural and functional dynamics underlying its regulation and inhibition is crucial for guiding therapeutic strategies targeting S1P signaling in disease.

It is well established that plasma membrane translocation and enzymatic activity of SK1 are enhanced by ERK1/2-mediated phosphorylation of Ser225. However, the structural mechanism by which this modification alters SK1 conformation remains unresolved^4,14^. Ser225 resides on the regulatory loop (R-loop), which is positioned opposite the C-terminal domain (CTD) β-sandwich core from the lipid-binding site. A longstanding question is how phosphorylation of this loop enhances membrane recruitment—required for sphingosine (Sph) phosphorylation to S1P—and catalytic activity. Notably, Ser225 is solvent-exposed in all available structures^15,16^. The R-loop tip packs against the N-terminal domain (NTD), stabilized by Asp235, which inserts into a pocket formed by basic residues (His156, Arg162, His355). It is postulated that phosphorylation of Ser225 displaces the R-loop from its NTD interaction, thereby increasing protein flexibility^4^. The functionally important 20 C-terminal residues of SK1 are missing from current crystal structures^15,16^. Truncation beyond residue 363 renders SK1 constitutively active and unresponsive to phorbol 12-myristate 13-acetate (PMA)^18^, while also enhancing membrane localization independently of Ser225 phosphorylation.

The lipid-binding loop 1 (LBL-1; Figs. 1a and 2a, purple segment) plays a key role in membrane interaction^4,19^. A hydrophobic patch containing Leu194, Phe197, and Leu198 (Fig. 2d) mediates curvature-sensitive membrane binding, which is important in endocytosis^4,20,21^. Together with Lys27, Lys29, and Arg186, these residues serve as key determinants for specific and nonspecific interactions with anionic phospholipid-enriched membranes^19^. The mechanism of sphingosine entry into the substrate-binding pocket remains unclear^4^. However, structural data suggest that conformational changes involving helices 7 and 8 (LBL-1) regulate substrate access, forming a flap-like structure over sphingosine^15^. Moreover, LBL-1 binds to the calcium- and integrin-binding protein (CIB1) via Phe197 and Leu198, which is thought to mediate SK1 membrane translocation^4,22^.

**Figure 1.**
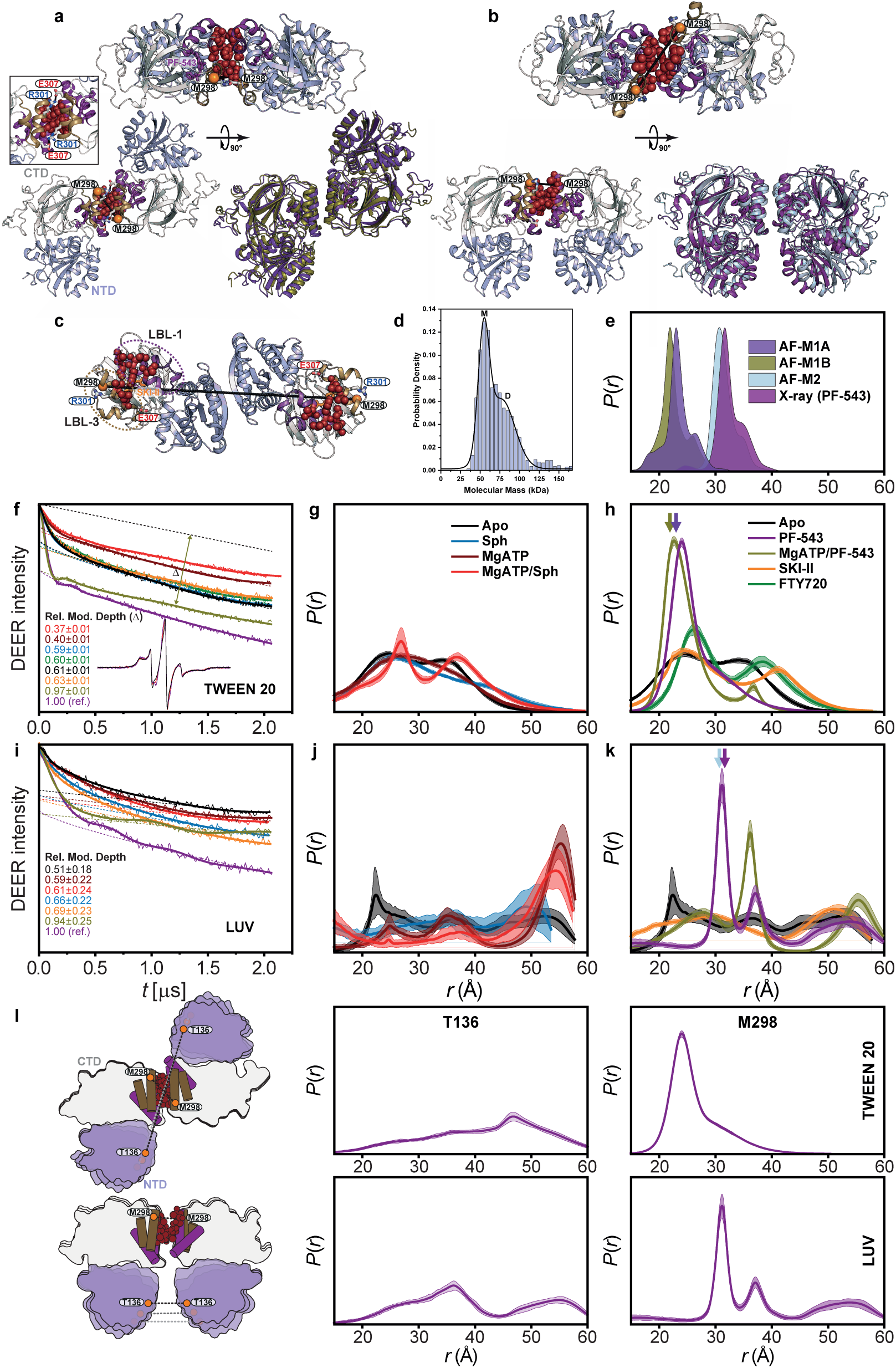
Potent inhibitors stabilize two distinct SK1 dimers via lipid-binding loops. (**a**) Cartoon representation of the PF-543–bound SK1 dimer model in Tween 20 (AF-M1/M1A, violet), with rotated monomers, aligned with the Mg^2+^ATP/PF-543 model (AF-M1B, olive). (**b**) PF-543–bound crystal structure (PDB 4V24, purple) aligned with the parallel, side-by-side dimer model (AF-M2, light blue). (**c**) Crystal structure of the SKI-II–bound SK1 dimer (PDB 3VZC, chains B and C), with the NTD, CTD, LBL-1, and LBL-3 shown in light blue, white, purple, and sand, respectively. Membrane-binding hydrophobic patches are shown as dark red spheres and spin-labeled M298 sites on LBL-3 as orange spheres. (**d**) Mass photometry of WT SK1 (100 nM) in detergent-free buffer reveals monomeric (M) and dimeric (D) populations. (**e**) Predicted distance distributions from the PF-543–bound crystal structure and dimer models. (**f**–**k**) Primary DEER traces with fits for singly labeled SK1 at M298 in the indicated ligand and inhibitor conditions, measured in Tween 20 (**f**–**h**) or liposomes (**i**–**k**), with corresponding distance distributions, *P*(*r*). The 95% confidence bands around the best-fit lines represent the estimated uncertainty in *P*(*r*) arising from fitting the primary DEER traces. Relative modulation depths (D) indicate dimerization relative to the PF-543–bound state. Catalytic complexes containing Mg^2+^ATP or Mg^2+^ATP/Sph exhibit limited, flexible dimerization, whereas PF-543 stabilizes well-defined dimers. (**l**) AF-M1 in detergent is stabilized by hydrophobic and electrostatic contacts, whereas AF-M2 in detergent-free buffer or with liposomes corresponds to the PF-543–bound crystal dimer. Thus, PF-543 stabilizes LBL-mediated SK1 dimers that block membrane binding and substrate access. Together, the structural models and DEER data support dimers s1tabilized primarily by CTD/LBL contacts, with dynamically oriented NTDs lacking persistent inter-protomer interactions.

**Figure 2.**
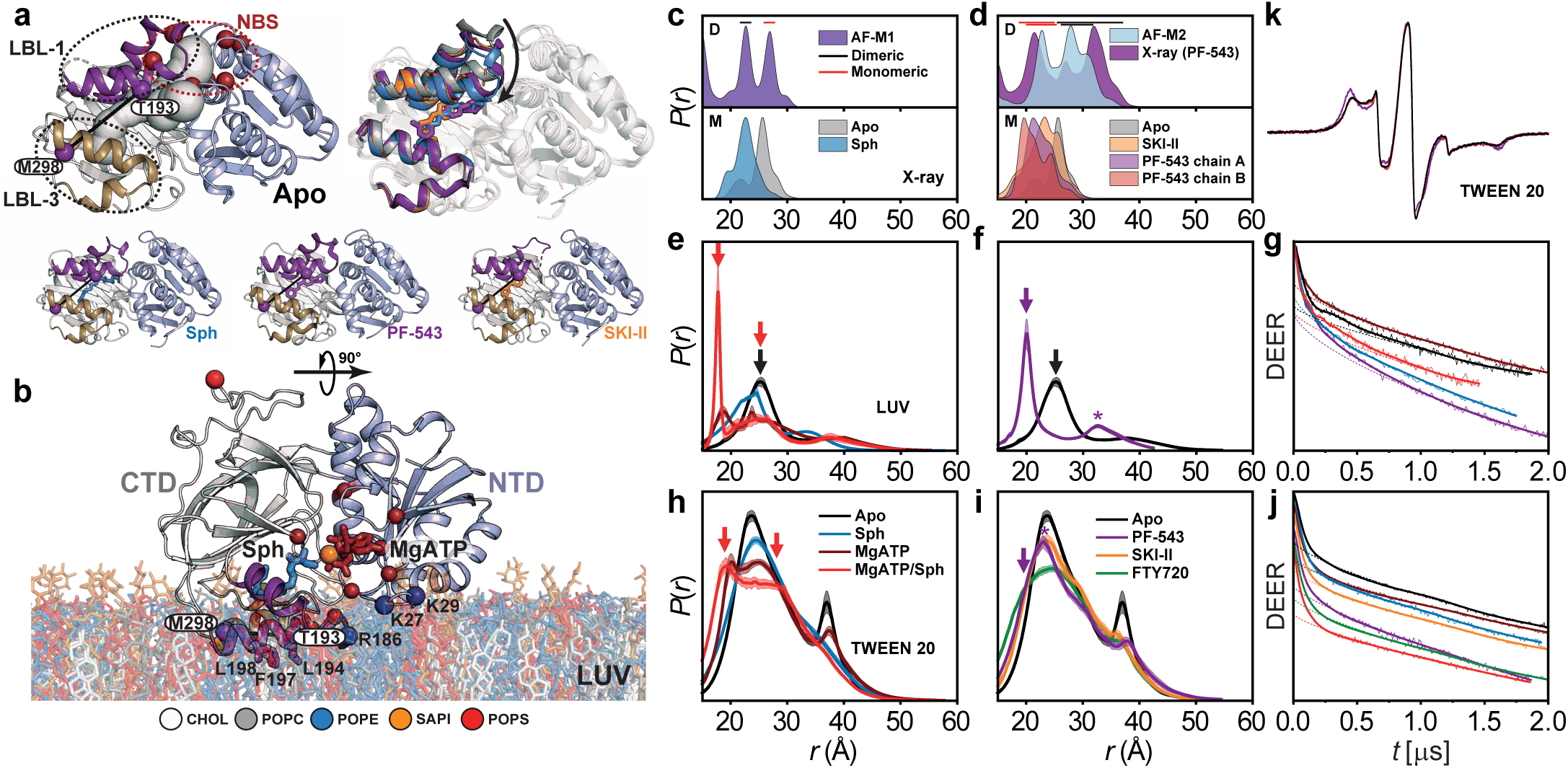
Ligand- and inhibitor-dependent conformational dynamics of SK1 lipid-binding loops. (**a**) Structural comparison of SK1 lipid-binding loops in the apo (PDB code 3VZB-C), sphingosine-bound (3VZB-A), PF-543-bound (4V24-A), and SKI-II-bound (3VZC-D) states, with the NTD, CTD, LBL-1, and LBL-3 colored light blue, white, purple, and sand, respectively. Spin label pair for DEER distance measurement (T193-M298, LBL gating) and Mg^2+^ATP-binding residues (NBS) are represented by purple and dark red spheres, respectively. Tunnels (white) calculated using MOLEonline 2.5 in the apo structure suggest that substrate entry from the membrane occurs through the LBL-1/LBL-3 gate, accessing the NBS occupied by Mg^2+^ATP. (**b**) Cartoon representation of Mg^2+^ATP/Sph-bound SK1 from our MD simulations, highlighting membrane-binding hydrophobic and electrostatic patches as purple sticks and dark blue spheres, respectively. (**c**,**d**) Upper panels: Predicted T193/M298 distance distributions from dimeric models and the PF-543–bound crystal structure, showing contributions from dimeric (black bar) and monomeric (red bar) states. Lower panels: Predicted monomeric distance distributions from crystal structures. (**e**–**j**) Distance distributions *P*(*r*) and primary DEER traces with fits with liposomes (LUV) (**e**–**g**) and in the presence of detergent (Tween 20) (**h**–**j**). Red, purple, and black arrows indicate structural intermediates preferentially populated in the catalytic complex (Mg^2+^ATP/Sph), PF-543-bound, and apo states, respectively. Purple asterisks indicate the dimeric contribution to the PF-543–bound distributions. (**k**) CW EPR spectra of the T193–M298 pair show no line-shape differences between the apo and Mg^2+^ATP/sphingosine states, but modest immobilization upon PF-543 binding, consistent with LBL-mediated dimerization that moderately restricts T193 on LBL-1. M298 mobility remains unchanged, consistent with its location at the start of LBL-3, away from the dimerization interface (Fig. 1f, inset). Sphingosine binding shifts the LBL-1/LBL-3 equilibrium toward a closed state, further stabilized by Mg^2+^ATP while still sampling both closed and open conformations, whereas PF-543 traps SK1 in a closed, low-dynamic state that blocks substrate access.

Some studies suggest that SK1 functions as a dimer^4,11,19,23^, but structural insights into how dimerization affects catalytic efficiency and regulation are limited. A putative dimeric assembly has been proposed based on existing structures^4,14,15^, with the N-terminal domains possibly forming the dimer interface. Nonetheless, the conformational ensemble and dynamics of SK1 dimers remain poorly characterized.

As a dynamic enzyme, understanding SK1’s conformational landscape is vital for the rational design of allosteric modulators^24–26^. Detailed structural knowledge of how weak and potent SK1 inhibitors, such as SKI-II and PF-543, engage the enzyme is still emerging^15–17^. PF-543 is a potent and selective SK1 inhibitor that acts via ATP-noncompetitive, substrate-competitive inhibition by mimicking sphingosine and occupying its binding pocket^16,27^. This prevents sphingosine access and phosphorylation. Key questions remain about how potent SK1 inhibitors, through orthosteric and allosteric effects, stabilize inactive conformations. Specifically, how do they reshape membrane-binding dynamics and how are these effects modulated by Mg^2+^ATP, given that most structural studies exclude ATP?

Despite considerable progress, several key mechanistic and structural questions remain. These include the conformational transitions involved in allosteric regulation and membrane association, as well as the functional implications of dimerization. The effect of dimerization on activity, substrate affinity, or allosteric modulation remains speculative. Moreover, SK1’s dynamic changes upon ligand binding are still insufficiently defined, posing a challenge to structure-based drug design. Although crystal structures of SK1 exist in both inhibitor- and substrate-bound states, high-resolution structures with both sphingosine and Mg^2+^ATP are lacking, and conformational shifts during catalysis remain uncharacterized. To address these gaps, we employed double electron–electron resonance (DEER; also known as PELDOR) spectroscopy^28–33^, AlphaFold modeling^34^, and all-atom molecular dynamics (MD) simulations^35–39^ to uncover new SK1 conformational states and elucidate the mechanistic basis of its regulation and inhibition— advancing our understanding of this important therapeutic target.

## Results

Mechanistic characterization of SK1 requires identifying its conformational states during substrate binding and catalysis, understanding their dynamic interconversion, and determining how post-translational modifications and inhibitors reshape this landscape to stabilize active or inactive states. To this end, DEER distance distributions were measured for SK1 in its apo form, bound to sphingosine with and without Mg^2+^ATP, bound to the sphingosine analog FTY720 (an immunomodulatory drug), and in complex with the inhibitors SKI-II and PF-543. Most measurements were conducted in the presence of 0.1% Tween 20, a nonionic detergent used in prior studies^15,19^. Additional DEER experiments for select pairs were performed in the presence of large unilamellar vesicles (LUVs) composed of a plasma membrane-mimetic composition.

### Functional integrity of SK1 mutants

To enable DEER spectroscopy, cysteine substitutions were introduced into a cysteine-less (CL) SK1 background. Functional integrity of the spin-labeled mutants was assessed using a fluorescence-based sphingosine kinase assay with 15-NBD-sphingosine^40^ and an ATPase activity assay^41^ (Supplementary Fig. 1). Most mutants retained activity comparable to, or at least 50% of, CL SK1. Notably, only D235N, which disrupts R-loop tethering, impaired sphingosine kinase activity while preserving ATPase activity (Supplementary Fig. 1b). Phosphomimetic (S225D) and phospho-null (S225A) substitutions at Ser225 caused only moderate reductions in activity relative to CL SK1 (Supplementary Figs. 1b and 1c), indicating that Ser225 modulates rather than dictates catalytic output. For direct comparison with WT SK1, S225D, D235N, and D235A were also introduced into the WT background. Gel-filtration profiles revealed increasing dimeric fractions in the order D235N > S225D > D235A > WT. Comparison of the monomeric and dimeric D235N fractions using the fluorometric SK1 assay showed that the dimer retained approximately half the activity of the monomer (Supplementary Fig. 1a, inset), supporting a potential regulatory role for SK1 dimerization. All kinetic measurements were performed using protein fractions that eluted as monomers (Supplementary Fig. 1).

### Lipid-binding loops mediate distinct SK1 dimerization modes

SK1 dimerization has been demonstrated in cells using immunoprecipitation assays^23^. Structural data also reveal two distinct potential dimerization interfaces—mediated by either the CTD or NTD domains—in SK1 bound to inhibitors PF-543 (PDB code 4V24; Fig. 1b)^16^ and SKI-II (PDB code 3VZC; Fig. 1c)^4,14,15^, respectively. Notably, in the PF-543-bound structure (Fig. 1b), hydrophobic residues on LBL-1 involved in membrane binding (Leu194, Phe197, Leu198) mediate dimerization, whereas in the SKI-II-bound structure (Fig. 1c), these same residues remain exposed and aligned for membrane interaction. However, these putative dimer interfaces have yet to be confirmed by biophysical methods and require further investigation. Mass photometry in detergent- and lipid-free buffer revealed SK1 dimerization (Fig. 1d), consistent with native mass spectrometry showing two distinct charge-state envelopes corresponding to ∼45 kDa and ∼90 kDa species (Supplementary Fig. 2a). To probe the quaternary structure of the SK1 dimer in both detergent (Figs. 1a and 1f–1h) and detergent-free liposome states (Figs. 1b and 1i–1k), we employed DEER spectroscopy with singly labeled SK1 protomers. Using the M298 site on LBL-3, we observed that under all ligand-bound conditions during substrate binding and catalysis, SK1 can adopt a highly flexible dimeric conformation to different extents, as reflected by broad distance distributions and large conformational ensembles (Figs. 1g and 1j), and the modulation depth (Δ) of DEER traces (Figs. 1f and 1i). Higher modulation depth indicates a higher concentration of interacting spin pairs, which is essential for quantifying monomer versus dimer populations in a mixture^28,42–44^. With consistent spin-labeling efficiencies (95–100%; e.g., 100% for M298 with 100±3 µM protein and 101±2 µM spin concentrations) and instrumental parameters, Δ reliably reflects the dimeric population^43^. In detergent, SK1 bound to Mg^2+^ATP or Mg^2+^ATP/sphingosine shows the lowest propensity for dimerization. By contrast, the potent inhibitor PF-543, with or without Mg^2+^ATP, markedly enhances dimerization, as evidenced by increased modulation depths (Figs. 1f and 1i) and pronounced oscillations in the primary DEER traces, indicating stabilization of distinct structured dimeric conformations (Figs. 1h and 1k). The observed distance populations correspond to two major classes of dimeric configurations predicted by AlphaFold (Figs. 1a, 1b, and 1e), both stabilized by interactions between lipid-binding loops 1 and 3, but differing in monomer orientation. In the first arrangement, observed with Tween 20 (AF-M1A and AF-M1B, Fig. 1a), the monomers are rotated relative to each other, and the LBL-1/LBL-3 interface buries ∼700 Å^2^ of solvent-accessible surface area per protomer. This interface is predominantly hydrophobic (LBL-1: Leu194, Phe197, Leu198, Leu200, Ala201; LBL-3: Leu300, Leu304), flanked by reciprocal hydrogen bonds between Arg301 and Leu304, and two salt bridges between Arg301 and Glu307 residues (Fig. 1a, inset), which act as strong “electrostatic clamps” stabilizing the interface. In contrast, the second arrangement (AF-M2, Fig. 1b), stabilized in detergent-free conditions, adopts a parallel, side-by-side orientation. Interestingly, this configuration also appears in the PF-543-bound crystal structure of SK1 solved in the absence of detergent, and may therefore represent a physiologically relevant dimer. Here, disruption of the Arg301–Glu307 ion pairs weakens the interface, yielding a more flexible dimer. Thus, the AF-M1 dimer appears more stable due to its combined polar and non-polar packing, compared with the AF-M2 and PF-543-bound crystal dimer that lack these salt bridges. Consistent with increased conformational flexibility, independent non-parametric and model-based DEER analyses converged on broader distance distributions for PF-543-bound, detergent-free samples (Fig. 1k and Supplementary Fig. 3) relative to those in Tween 20 (Fig. 1h). These analyses further reveal weak, heterogeneous dimerization across apo and catalytic conditions, including Mg^2+^ATP and Mg^2+^ATP/Sph, reflected in broad DEER ensembles (Fig. 1 and Supplementary Fig. 3). Importantly, the NTDs lack direct contacts in both dimer classes (Fig. 1l), implying a dynamic orientation between them. DEER measurements at NTD residue T136 revealed broad inter-NTD distance distributions in both Tween 20 and LUVs, consistent with substantial conformational heterogeneity (Fig. 1l and Supplementary Fig. 2). In the SKI-II-bound structure, the dimer places M298 residues ∼10 nm apart (Fig. 1c), a distance not detected in our DEER measurements. As previously proposed^4,19,20^, factors such as membrane curvature may stabilize this conformation, warranting further investigation.

Together, these findings reveal that PF-543 as a potent inhibitor stabilizes LBL-mediated dimeric structures, highlighting a novel inhibition mechanism in which membrane interaction through LBLs and substrate access are hindered.

### Dynamic lipid-binding loops regulate sphingosine entry, with PF-543 locking a closed conformation

The mechanism by which sphingosine accesses its binding site in SK1 has remained unclear. Crystal structures (PDB code 3VZB) show sphingosine enclosed within the CTD, implying full lipid extraction (Figs. 2a and 2b). The only apparent entry is a narrow path through the ATP-binding site, which would require tail-first tunneling past polar residues. A more plausible mechanism involves dynamic opening of LBL-1, which form a flap over the binding site (Fig. 2b). Structural comparisons between apo and ligand-bound states suggest that LBL-1 rotates outward to expose inner hydrophobic surfaces and the β-sandwich core (Fig. 2a)^4,15^. However, crystal packing may restrict this movement, necessitating experimental validation. To investigate LBL conformational ensemble during substrate and inhibitor binding, we monitored the T193–M298 distance (LBL-1 to LBL-3). CW EPR spectra of this pair were unchanged between the apo and Mg^2+^ATP/sphingosine states but showed modest immobilization upon PF-543 binding, consistent with LBL-mediated dimerization that partially restricts T193 on LBL-1 (Fig. 2k)^45^. In contrast, M298 mobility remained unchanged, consistent with its position at the beginning of LBL-3, distal to the dimerization interface (Fig. 1f, inset). Since SK1 only partially adopts highly dynamic dimeric structures under most conditions, whereas PF-543 stabilizes two distinct conformations (AF-M1 in Tween 20 and AF-M2 in LUVs, consistent with the PF-543 X-ray structure), we can integrate this knowledge with existing crystal structures to distinguish dimeric (black bar) and monomeric (red bar) contributions in the DEER distance distributions (Figs. 2c and 2d). Despite high labeling efficiency, the limited pump inversion efficiency and low modulation depths minimize multispin ghost contributions and their impact on the distance distributions^46,47^. In the apo state—especially with liposomes— measured distances matched the apo structure^15^, indicating an open LBL-1 (Fig. 2e and Supplementary Fig. 4). Sphingosine binding shifted the distribution toward shorter distances, particularly in liposomes, consistent with the more closed conformation captured in the sphingosine-bound crystal structure. Binding of Mg^2+^ATP/sphingosine stabilizes a closed state not captured in crystal structures or dimeric models (red arrow, Figs. 2e and 2h and Supplementary Fig. 4), while Mg^2+^ATP alone partially shifted the equilibrium (Figs. 2e and 2h). These findings support a model where transient LBL-1 opening enables substrate recognition at the membrane interface, followed by substrate extraction and LBL-1 closure (Fig. 2b). In both detergent and liposomes, ligand binding increases LBL dynamics, resulting in a broader conformational ensemble, particularly in the catalytic Mg^2+^ATP/sphingosine complex, which samples both closed and open states (Figs. 2e and 2h, red arrows). This flexibility likely facilitates phosphoryl transfer and S1P release. Sphingosine analog FTY720 and SKI-II induce similar LBL behavior (Fig. 2i), whereas PF-543 strongly favors the closed conformation and reduces LBL mobility, suggesting an inhibition mechanism that, in addition to LBL-mediated dimerization that hampers membrane binding, locks the loops in a closed state and blocks substrate access (Figs. 2f and 2i). Thus, LBL dynamics regulate sphingosine entry and S1P release, whereas potent inhibitors such as PF-543 mostly trap SK1 in a closed conformation (excluding dimeric contributions), blocking substrate access.

### Ser225 phosphorylation reconfigures the R-loop into a catalytic state, modulated by PF-543

Phosphorylation at Ser225, a key regulator of SK1’s activity and membrane localization, induces conformational changes that remain incompletely characterized. Ser225 lies exposed within the R-loop, on the reverse side of the CTD β-sandwich, away from the lipid-binding site (Fig. 3b). The R-loop tip contacts NTD helices α3/α4 and is stabilized by Asp235, which inserts into a β-sandwich pocket formed by His156, Arg162, and His355 (Fig. 3b). It is proposed that phosphorylation displaces Asp235, altering R-loop–NTD interactions and aligning membrane-facing residues in the NTD and CTD (Fig. 3b, blue spheres) for optimal membrane binding^4,14^. To monitor the R-loop conformational ensemble, the S159-V234 distance pair reports on R-loop movement near Asp235 (Fig. 3 and Supplementary Fig. 5). The monomeric and dimeric distance components are clearly separated (Figs. 3c–3e). Across WT conditions, with minor variations, the dominant distance population corresponds to those observed in the apo and sphingosine-bound structures, indicating a stable Asp235 salt bridge critical for catalysis (Figs. 3f and 3g)^15^. Interestingly, the D235N mutation enhances R-loop dynamics and shifts the population toward longer distances, indicating salt bridge disruption (Figs. 3h and 3i). This effect is also evident in CW-EPR spectra compared with the WT protein^45^. The T99–A224 and S159–T222 pairs monitor changes near Ser225 (Fig. 3 and Supplementary Fig. 6), revealing two distinct conformations (States 1 and 2, Figs. 3j-3q). Interestingly, AlphaFold modeling of phosphorylated SK1 (Ser225^P^) predicts a conformation where Ser225^P^ forms a salt bridge with His156/Arg162, replacing Asp235, while the Asp235–His355 salt bridge remains intact (State 1, Fig. 3d, inset). Importantly, this configuration corresponds to the experimentally determined DEER ensemble and is most populated in in the Mg^2+^ATP-or Mg^2+^ATP/Sph-bound states (State 1, Figs. 3j, 3n, and 3o), whereas it is least populated in sphingosine-bound state. Consistent with previous studies^13,16^, our mass spectrometry analysis confirms partial Ser225 phosphorylation in our constructs (Fig. 3a), indicating potential conformational heterogeneity. Distance predictions from dimeric models with engaged Ser225^P^ (red, Figs. 3d and 3e) versus detached states show that the T99–A224 and S159–T222 distance pairs can distinguish between the two, with States 1 and 2 in DEER data representing the engaged and detached populations, respectively. For the S159–T222 pair, the phosphomimetic S225D mutant closely recapitulates WT behavior, with a more pronounced shift toward the engaged state under Mg^2+^ATP-bound conditions (Fig. 3l). In contrast, the phospho-null S225A mutant markedly reduces State 1 and yields more overlapping distance populations (Fig. 3m).

**Figure 3.**
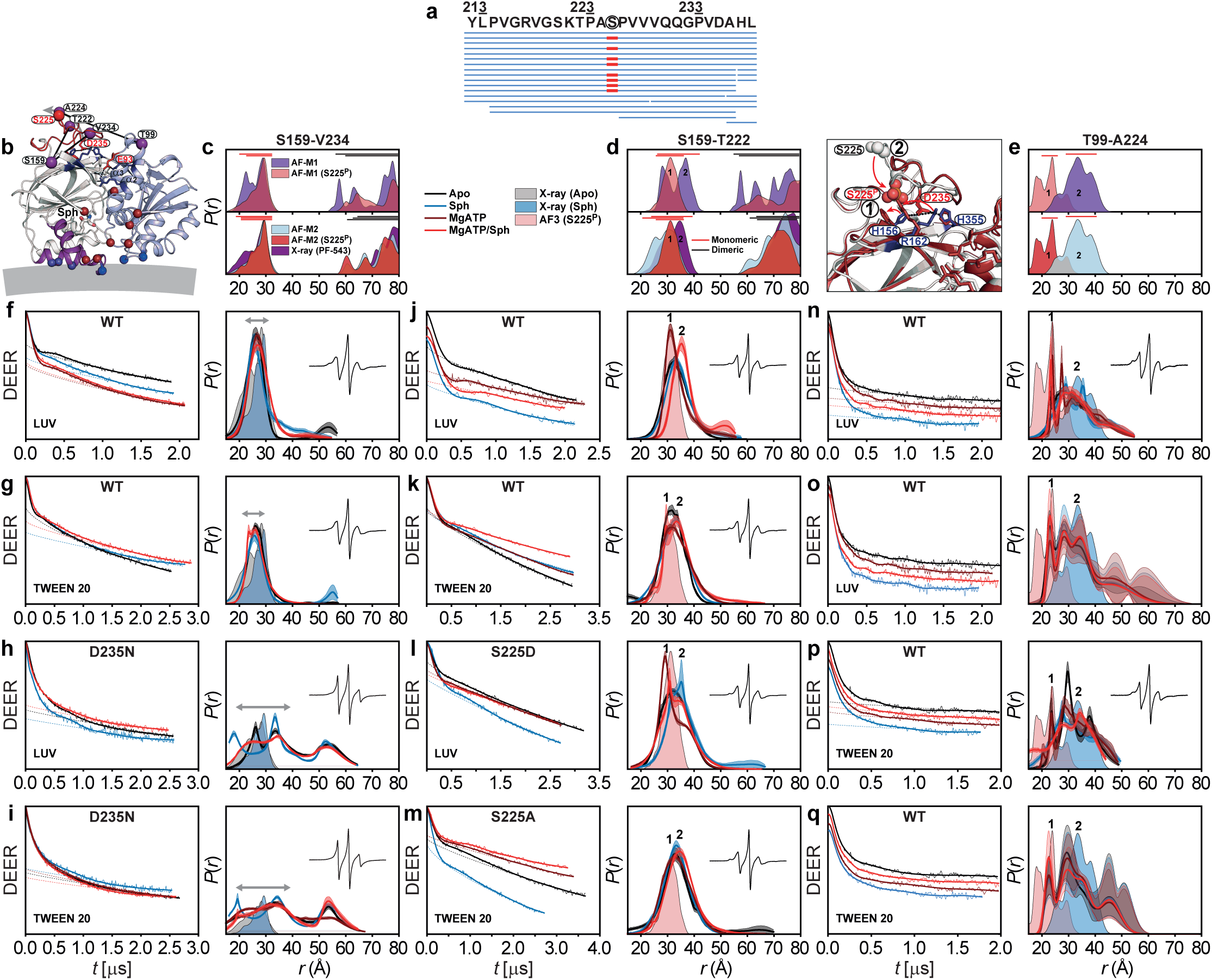
Ser225 phosphoryIation reshapes the R-Ioop into a cataIyticaIIy competent state. (**a**) Mass spectrometry identified multiple overlapping peptides spanning Ser225, several containing phosphorylation (red), confirming site-specific phosphorylation in a subset of the protein population. (**b**) DEER distance pairs S159–T222, S159–V234, and T99–A224 (purple spheres) mapped onto the sphingosine-bound structure, with the NTD, CTD, R-loop, and LBL-1 shown in light blue, white, red, and purple, respectively. Functional residues, including S225 and D235, are labeled in red; R-loop salt-bridge networks are shown as sticks, and Mg^2+^ATP-binding and membrane-interacting residues as dark red and blue spheres. (**c**–**e**) Predicted distance distributions for dimeric models (AF-M1 and AF-M2) with or without engaged Ser225^P^ (States **1** and **2**, inset), together with the PF-543–bound crystal structure, reveal distinct monomeric (red bar) and dimeric (black bar) contributions. AF3 modeling predicts that Ser225^P^ forms new salt bridges with His156/Arg162 while Asp235 retains contacts with His355, consistent with the experimental ensemble. T99–A224 and S159–T222 distinguish engaged Ser225^P^ (State **1**) from the detached state (State **2**). (**f**,**g**) DEER measurements of S159–V234 indicate a stable Asp235–His355 interaction in WT SK1. (**h**,**i**) D235N disrupts this pocket and shifts the ensemble toward longer distances, consistent with increased flexibility and CW EPR spectra (insets). Double-headed gray arrows indicate the dynamic range of R-loop conformations around Asp235. (**j**–**q**) DEER distance distributions probe conformational ensembles near partially phosphorylated Ser225: S159–T222 in WT (**j**,**k**), the phosphomimetic S225D (**I**), and phospho-null S225A mutation (**m**); and T99–A224 pair analyzed by model-free neural network approach (**n**,**p**) and global Gaussian fitting (**o**,**q**). Two conformations emerge: State 1, favored by Mg^2+^ATP or Mg^2+^ATP/Sph, reflects engaged Ser225^P^ interactions, whereas State 2 predominates in sphingosine-bound SK1. S225D resembles WT, supporting a model in which Ser225 phosphorylation promotes a catalytically competent R-loop configuration modulated by ligand binding. Predicted distributions from the apo and sphingosine-bound crystal structures and the AF3 model with engaged Ser225^P^ are shaded light gray, blue, and red, respectively.

To further investigate the rearrangement of salt bridges between the R-loop and the strand pair connecting the NTD to the CTD upon Ser225 phosphorylation, we conducted all-atom equilibrium MD simulations in the presence and absence of ligands and R-loop phosphorylation (Fig. 4 and Supplementary Figs. 7 and 8). Across all systems, the root mean square deviation (RMSD) stabilizes at ∼2–3 Å, indicating that SK1 maintains overall conformational stability in the membrane-bound state (Supplementary Figs. 7a and 7b). Apo systems, independent of Ser225 phosphorylation, exhibit slightly higher flexibility than ligand-bound systems, whereas Mg^2+^ATP/Sph-bound systems display reduced RMSD variability, consistent with ligand-induced stabilization. The Mg^2+^ATP/Sph system with engaged Ser225^P^ converges most rapidly and exhibits the highest stability. Root mean-square fluctuation (RMSF) analysis shows that the regulatory loop is highly flexible in apo systems but becomes progressively restrained upon Mg^2+^ATP/sphingosine binding and Ser225 phosphorylation (Supplementary Fig. 7c). Collectively, these results indicate that Mg^2+^ATP/Sph binding and Ser225 phosphorylation act in concert to allosterically reinforce SK1 conformational stability. Substrate binding alone does not reconfigure R-loop salt bridges: the stable Asp235–His355/His156 interactions remain, while the weaker, transient Asp235–Arg162 contact persists (Mg^2+^ATP/Sph panels, Fig. 4c). In this state, Ser225 is fully disengaged from these basic residues (Fig. 4b and Supplementary Fig. 8b). MD simulations further show that Ser225 phosphorylation stabilizes salt bridges with His156/Arg162 and reconfigures the R-loop, especially in the presence of substrates (Mg^2+^ATP/Sph(Ser225^P^) panels in Fig. 4b and Supplementary Fig. 8b and Supplementary Video 1). Thus, consistent with the DEER measurements (Figs. 3j, 3l, 3n, 3o), substrate binding mutually and allosterically stabilizes this R-loop conformation. Although phosphorylation alone can form these bridges, they are less stable without substrates (Supplementary Fig. 8b and Supplementary Video 2). Asp235 consistently maintains interaction with His355, but its salt bridge with His156 weakens upon phosphorylation (Fig. 4c and Supplementary Fig. 8c). These results underscore Ser225 phosphorylation as a key allosteric switch for adopting a catalytically competent conformation.

**Figure 4.**
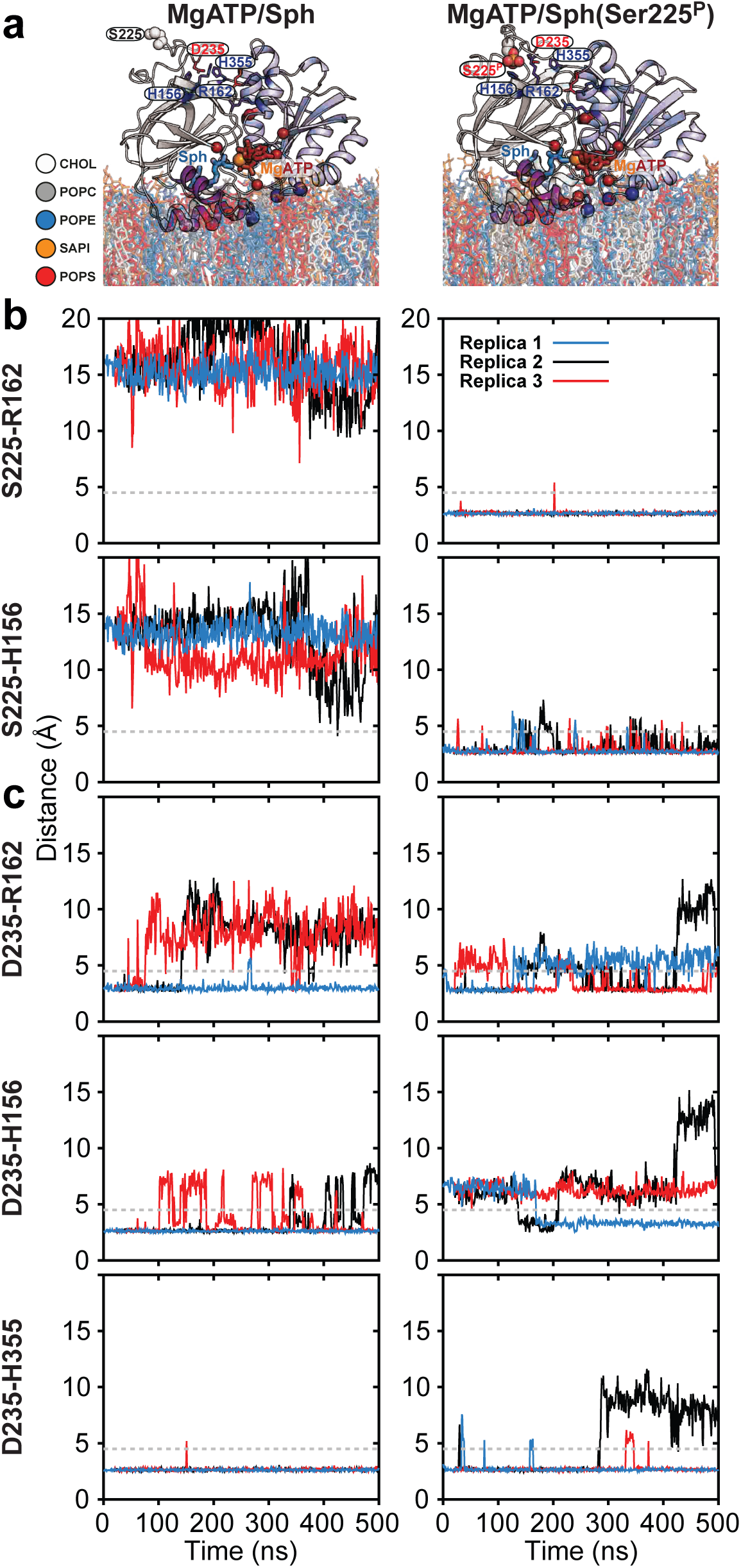
Substrate binding mutually and allosterically stabilizes the reconfigured R-loop conformation following Ser225 phosphorylation. (**a**) Two MD simulation systems of ligand-bound SK1 bound to plasma membrane lipid bilayers, with and without R-loop phosphorylation, were each performed in three independent replicates (Supplementary Fig. 7). (**b**,**c**) Time series of the distances between basic residue side chains and either Ser225/Ser225^P^ or Asp235 during simulations, used to monitor salt bridge formation between the R-loop and the strand pair connecting the NTD to the CTD. Phosphorylation of Ser225 stabilizes salt bridges to His156 and Arg162, reconfiguring the R-loop. Binding of substrates (Mg^2+^ATP and Sph) stabilizes this R-loop conformation. Regardless of the condition, Asp235 consistently maintains a tight interaction with His355, while the salt bridge with His156 is less stable and becomes disrupted upon Ser225 phosphorylation.

FTY720 and inhibitors modulate R-loop conformations (Fig. 5). Beyond enhanced dimerization, the S159–T222 and T99–A224 pairs show a markedly reduced population of the catalytically competent, engaged Ser225^P^ state (State 1), particularly in the presence of PF-543 (Fig. 5 and Supplementary Fig. 6). Phosphomimetic S225D and phospho-null S225A mutations behave similarly to WT Additional residues, such as Asn89 and Glu93 (Fig. 3b)^4^, may also contribute to phosphorylation-induced membrane targeting. Asn89, in particular, forms potentially stabilizing cross-domain hydrogen bonds that may help align the NTD and CTD for optimal membrane binding.

**Figure 5.**
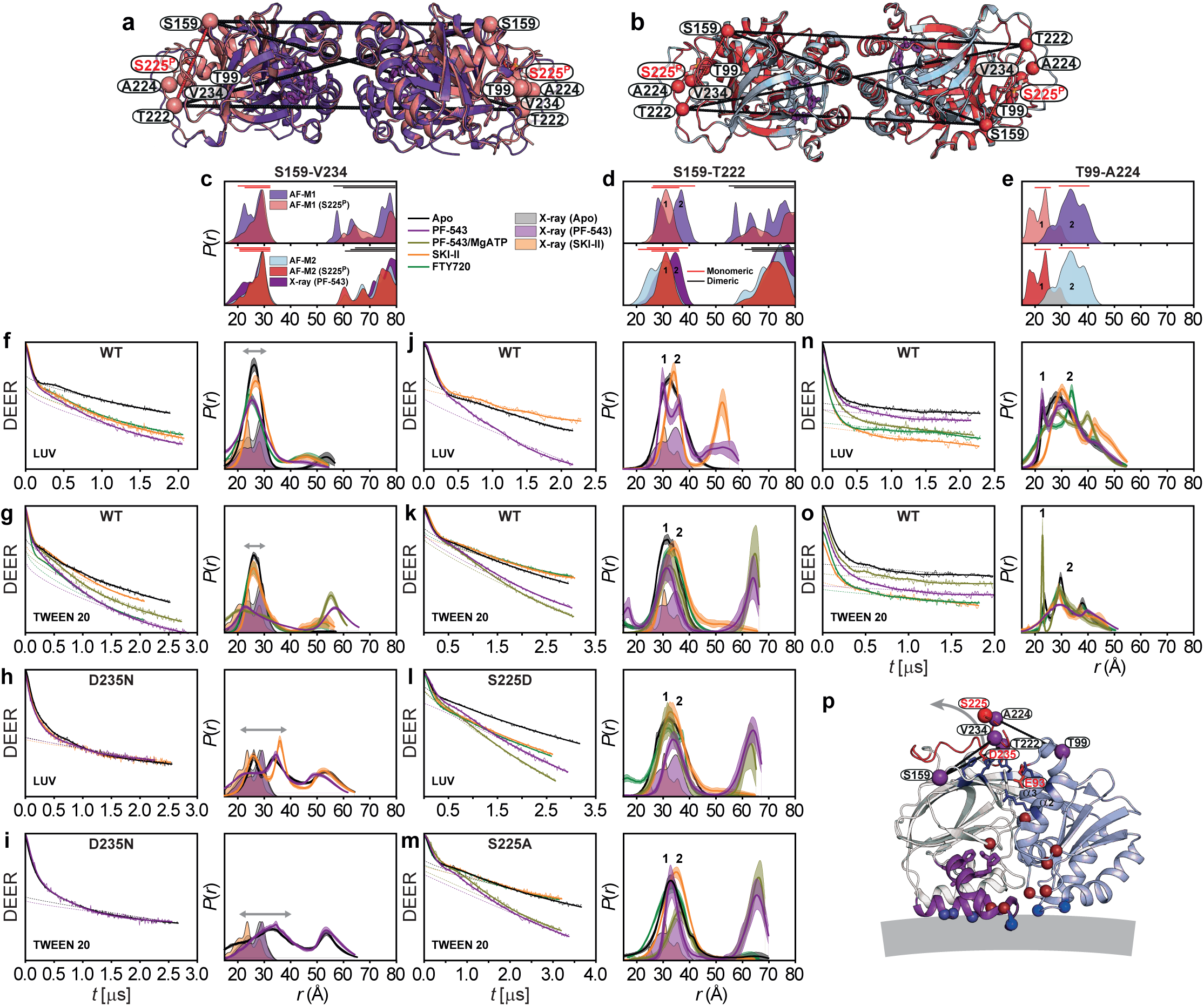
Inhibitors reshape R-Ioop conformationaI ensembIes. (**a**,**b**) PF-543–stabilized dimeric models showing the DEER pairs S159–V234, S159–T222, and T99–A224 used to probe R-loop conformations. For clarity, only dimeric (black) and monomeric (red) distances are shown for S159–T222 in models with engaged Ser225^P^. (**c**–**e**) Predicted dis1tance distributions from dimeric models and the PF-543–bound crystal structure. (**f**–**o**) DEER traces and corresponding distance distributions, *P*(*r*), probing R-loop conformations in the presence of inhibitors and the sphingosine analog FTY720 for WT SK1 and the D235N, S225D, and S225A mutants in detergent (Tween 20) and liposomes (LUV). Confidence bands around the best fits represent the estimated uncertainty in *P*(*r*). Grey arrows indicate the dynamic range of R-loop conformations around Asp235. States 1 and 2 (**d**,**e**,**j**–**o**) correspond to engaged Ser225^P^ and detached conformations, respectively. Predicted distributions from the apo, PF-543–bound, and SKI-II–bound crystal structures are shaded light grey, purple, and orange, respectively. (**p**) DEER pairs T99–A224, S159–T222 and S159–V234 (purple spheres) mapped onto the PF-543–bound structure, with the NTD, CTD, and LBL-1 shown in light blue, white, and purple, respectively. Functional residues, including S225 and D235, are labeled in red; R-loop salt-bridge networks are shown as sticks, and Mg^2+^ATP-binding and membrane-interacting residues as dark red and blue spheres. Together, these data indicate that inhibitors reshape the R-loop conformational ensemble beyond their effects on dimerization, generally shifting the ensemble away from the catalytically competent, engaged Ser225^P^ state (State **1**) toward detached conformations (State **2**).

### SK1 adopts a novel catalytic conformation inhibited by PF-543

We introduced interdomain distance pairs between the NTD and CTD to investigate their relative orientation and dynamics, which are regulated by ligand and inhibitor binding, as well as S1P synthesis (Fig. 6 and Supplementary Fig. 9). Sphingosine, its analogs, and inhibitors bind to the CTD, while Mg^2+^ATP binds to the NTD at the domain interface (Fig. 6a). Except for the T136-T193 pair, the catalytic complex with Mg^2+^ATP/sphingosine is well-structured (Figs. 6b, 6c, 6k, and 6l and Supplementary Fig. 9). The protein remains highly dynamic in the apo and sphingosine-bound states. The catalytic complex, which has never been structurally studied before, adopts a conformation distinct from the apo and sphingosine-bound crystal structures (Fig. 6, red asterisk). Interestingly, even after separating potential dimeric contributions to the DEER distance distributions (Figs. 6g and 6j), the T136-T193 pair, which includes the catalytic LBL-1 region, remains highly dynamic in all states, including the catalytic complex (Figs. 6f and 6i and Supplementary Fig. 9), and deviates significantly from the apo and sphingosine-bound crystal structures. However, such conformational flexibility is absent in the PF-543–bound state, which instead favors shorter-distance intermediates, excluding dimeric contributions, regardless of Mg^2+^ATP (Figs. 6f and 6i). For the T136-T193 pair, this domain configuration aligns with the PF-543-bound crystal structure, suggesting that PF-543 stabilizes a non-catalytic conformation, unlike sphingosine or SKI-II, regardless of Mg^2+^ATP binding. In contrast, the catalytic complex maintains LBL-1 flexibility (Figs. 2e, 2h, 6e, and 6f), which likely supports S1P production and release. PF-543 partially stabilizes the catalytic conformation at interdomain distance pair T136–M298, with one site located on the LBL-3 (Fig. 6c, purple asterisk), a trend also observed for the low-activity T136–A305 mutant (Fig. 6l). MD simulations reveal distinct dynamic behaviors of LBL-1 and LBL-3 across states (Supplementary Fig. 7c). These findings provide additional structural insights into the inhibition mechanism of PF-543.

**Figure 6.**
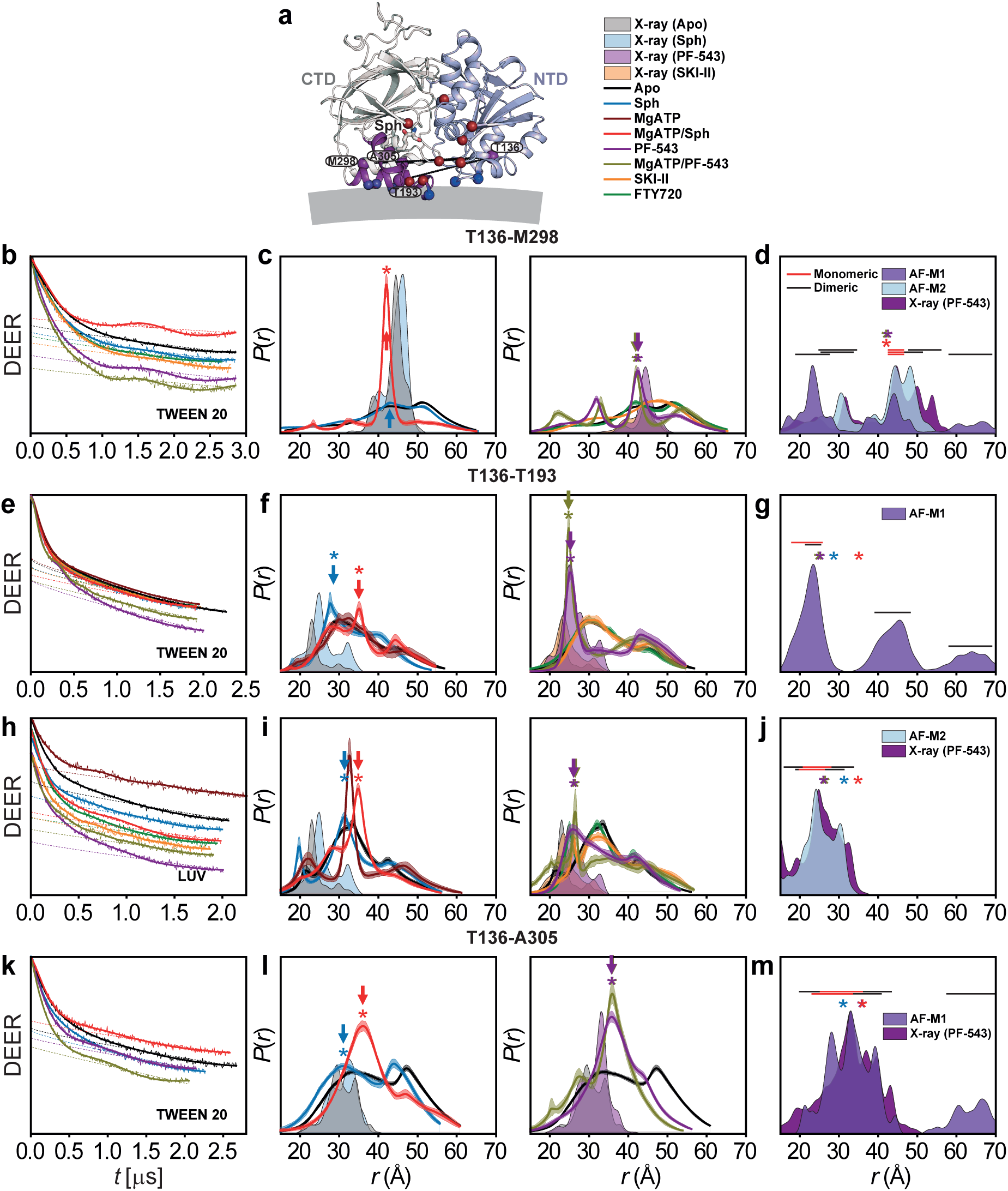
SK1 adopts a noveI cataIytic conformation with a dynamic LBL-1, inhibited by PF-543. (**a**) Interdomain DEER distance pairs (T136-M298, T136-T193, T136-A305, purple spheres) probing the catalytic core mapped onto the sphingosine-bound structure, with the NTD, CTD, and LBL-1 shown in light blue, white, and purple, respectively. Mg^2+^ATP-binding and membrane-interacting residues are depicted as dark red and blue spheres. (**b**–**m**) Primary DEER traces and distance distributions *P*(*r*) for the indicated pairs in detergent (Tween 20) and liposomes (LUV), compared with predictions from crystal structures and AF models. Red, blue, purple, and olive arrows and asterisks denote intermediates populated in the catalytic Mg^2+^ATP/sphingosine complex, sphingosine-bound, PF-543–bound, and Mg^2+^ATP/PF-543–bound states, respectively. For most pairs, the catalytic complex with Mg^2+^ATP/sphingosine adopts a distinct conformation not observed in apo or sphingosine-bound structures (red asterisks), consistent with a previously uncharacterized catalytic state. While most distance pairs stabilize in this complex, the T136–T193 pair probing the catalytic LBL-1 region remains highly dynamic, suggesting flexibility important for S1P synthesis and release. This flexibility is lost in the PF-543–bound state, which instead favors shorter-distance intermediates aligned with the PF-543 crystal structure, independent of Mg^2+^ATP binding, indicating stabilization of a non-catalytic conformation. PF-543 also partially stabilizes the catalytic configuration at the T136–M298 pair, a trend mirrored by the low-activity T136–A305 mutant. These results show that PF-543 selectively suppresses LBL-1 flexibility and traps SK1 in a non-productive state, providing a structural basis for its inhibitory mechanism.

### The C-terminal tail displays distinct proximal–distal states

The C-terminal tail (residues 365–384), absent from all crystal structures, harbors known protein interaction sites and plays a key regulatory role (Fig. 7a)^4,14,18^. Interestingly, its interaction with the long, twisted strand pair that connects the NTD and CTD—implicated in R-loop anchoring—has been postulated (see Discussion)^14^. In this context, Ser225 phosphorylation-induced structural transitions in the R-loop may promote C-terminal tail release, facilitating membrane targeting. CW EPR spectrum of spin-labeled Val366 at the tail’s N-terminus showed intermediate spin label motion^45^, indicating contact with the core protein (Fig. 7b)^19^. In contrast, the Trp372 spectrum exhibited sharper, narrower lines, suggesting a more flexible conformation and greater mobility for this site and subsequent residues. Given Ser225 phosphorylation in our constructs, this supports a model in which the tail is released from its potential tethering to the core protein. DEER measurements reveal broader distance distributions for T136–W372 pair compared with T136–V366 (Figs. 7c–7h and Supplementary Fig. 10). For the T136–V366 pair, catalytic conditions maintained a predominantly short-distance population while producing condition-dependent redistribution within the ensemble (Fig. 7d). Inhibitor-bound states similarly preserved association of the proximal C-terminal tail with the protein core but altered the underlying conformational distribution (Fig. 7e). By contrast, T136–W372 exhibited broader and more overlapping distance distributions across conditions, consistent with greater conformational heterogeneity and weaker coupling of the distal tail to the core (Figs. 7f–7h and Supplementary Fig. 10). Measurements of singly labeled V366 and W372 further revealed broad, multimodal intermolecular tail–tail distance distributions (Figs. 7i–7n). Within dimeric assemblies, the proximal tail showed modest ligand-dependent reorganization, whereas the distal region remained substantially more heterogeneous. Collectively, these observations support a segmentally organized C-terminal tail in which a relatively core-associated, condition-responsive proximal segment transitions into a highly dynamic distal region. Both catalytic and inhibitory states reshape this conformational ensemble, suggesting a potential coupling between R-loop regulation and C-terminal-tail organization.

**Figure 7.**
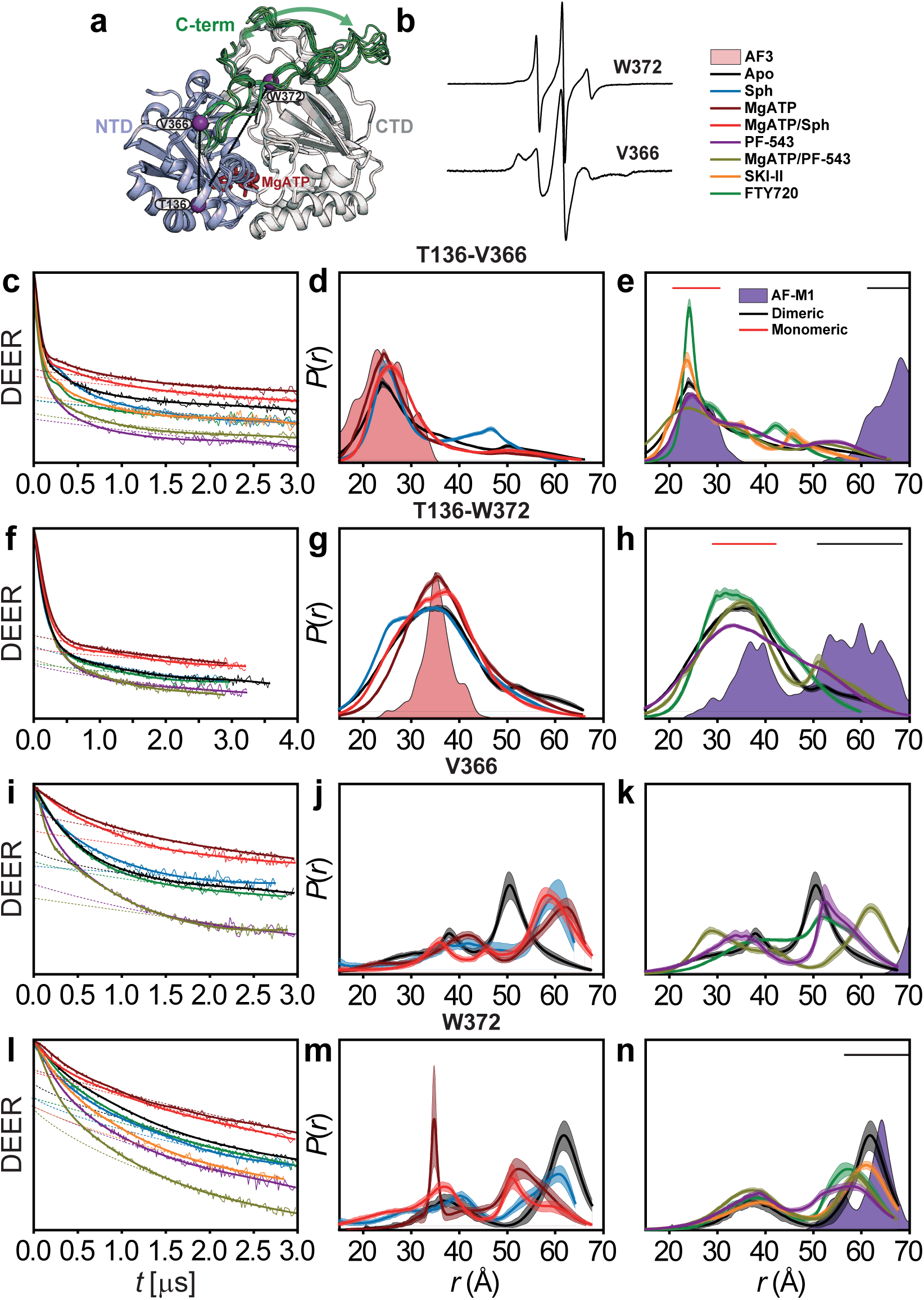
The C-terminaI taiI comprises a core-associated proximaI region and a dynamic distaI segment. The C-terminal tail (residues 365–384), absent from available crystal structures but implicated in regulatory interactions, shows distinct segmental dynamics by CW EPR and DEER. (**a**) An ensemble of AlphaFold3 models of Mg^2+^ATP-bound SK1 with an intact C-terminal tail (green) illustrates its conformational heterogeneity. DEER pairs T136–V366 and T136–W372 (purple spheres) are mapped onto a representative model consistent with the major experimental distance populations. (**b**) CW EPR spectra show intermediate mobility at V366, consistent with partial core association, and greater mobility at W372, indicating increased distal-tail flexibility. (**c**–**e**) Primary DEER traces and distance distributions, *P*(*r*), for T136–V366. Catalytic conditions (**d**) retain a dominant short-distance population with condition-dependent redistribution, whereas inhibitor-bound states (**e**) preserve proximal core association while reshaping the ensemble. The AlphaFold3-predicted distribution is shaded light red. (**f**–**h**) Corresponding T136–W372 measurements show broader, more overlapping distributions across conditions, consistent with greater heterogeneity and weaker distal-tail coupling to the protein core (Supplementary Fig. 10). (**i**–**n**) DEER measurements of singly labeled V366 and W372 reveal broad, multimodal intermolecular tail–tail distance distributions, with modest ligand-dependent reorganization of the proximal tail within dimeric assemblies and greater heterogeneity distally. Together, these data support a segmentally organized C-terminal tail, with a relatively core-associated, condition-sensitive proximal region transitioning to a highly dynamic distal segment. Catalytic and inhibitory states remodel this ensemble, potentially coupling R-loop regulation to C-terminal-tail organization.

## Discussion

Using an integrated spectroscopic and computational approach, this study reveals new mechanistic insights into SK1, uncovering a dynamic, multilayered regulatory mechanism in which structural flexibility governs both catalysis and inhibition. A key breakthrough is the discovery that potent SK1 inhibitors not only stabilize non-catalytic conformations but also drive dimerization via lipid-binding loops (LBLs), unveiling a novel inhibitory mechanism that blocks membrane binding and substrate access. Notably, LBL-mediated dimerization may also occur under cellular conditions, serving as both an inhibitory and regulatory mechanism. Another key finding is that LBL dynamics gate sphingosine entry into the active site, with the potent inhibitor PF-543 locking the LBLs in a closed, less dynamic conformation that blocks ligand exchange (Fig. 2f). A transformative finding is that phosphorylation of Ser225 may reconfigure the R-loop and the overall SK1 structure into a catalytically competent state. Comprehensive MD simulations reveal a reshuffling of salt bridges involving positively charged residues on the strand pairs connecting the two SK1 lobes—potentially facilitating membrane engagement (Figs. 4 and 8 and Supplementary Fig. 8) and release of the C-terminal tail. These simulations uncover phosphorylation-and ligand-dependent shifts in the balance between electrostatic and hydrophobic interactions in membrane engagement, suggesting a dynamic mechanism by which SK1 senses and responds to the membrane environment (Fig. 8). The catalytically active SK1 adopts a previously uncharacterized conformation, distinct from known apo or ligand-bound structures, featuring a highly dynamic LBL-1 region that is sensitive to PF-543, which stabilizes a non-catalytic state (Figs. 1 and 6).

**Figure 8.**
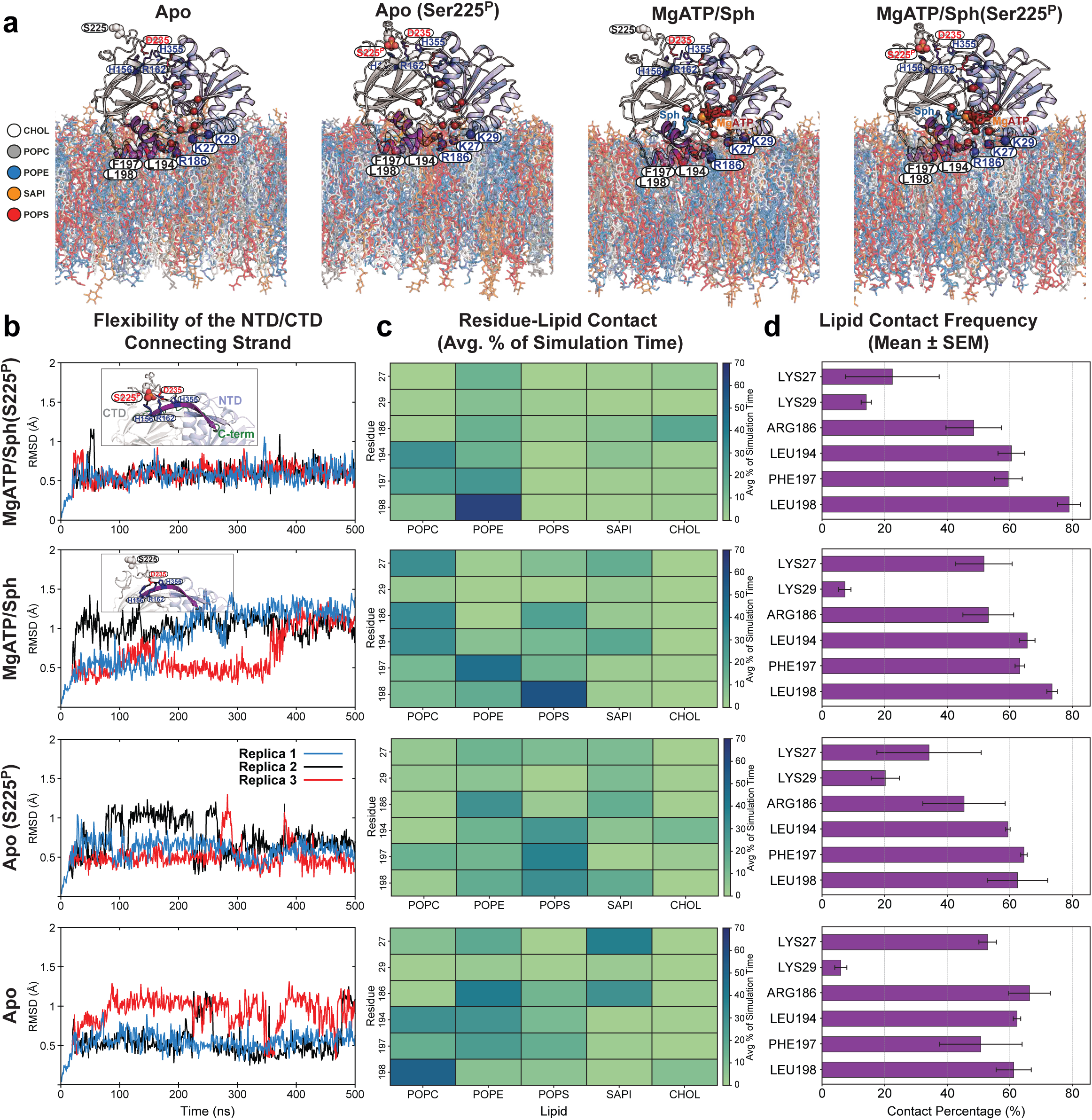
Phosphorylation-dependent modulation of SK1 connecting strand flexibility and membrane engagement. (**a**) Representative snapshots of four MD simulation systems, each performed in three independent replicates. Membrane-interacting residues are shown as dark blue (basic) and purple (hydrophobic) spheres. (**b**) Flexibility of the NTD/CTD connecting strand (residues 350-362, shown in purple in the inset) in ligand-bound SK1 is significantly reduced upon Ser225 phosphorylation, likely due to the formation of stable salt bridges (i.e., Ser225^P^:Arg162, Ser225^P^:His156, and Asp235:His355; see Supplementary Fig. 7). (**c**) Lipid interaction specificity of membrane-interfacing residues, plotted as the average contact percentage of total simulation time, reveals distinct interaction patterns for phosphorylated versus non-phosphorylated SK1. Electrostatic interactions between basic residues and anionic lipids are significantly diminished in the Mg^2+^ATP/Sph-bound phosphorylated protein, while hydrophobic interactions increase. In the apo state, Arg186 and Lys27 preferentially interact with SAPI over POPS, with Arg186 also contacting POPE. In Apo(Ser225^P^) state, the hydrophobic patch shows increased contact with POPS. (**d**) Overall, the lipid contact frequency of the hydrophobic patch and Arg186 remains high across conditions, whereas Lys27 lipid interactions are significantly reduced upon Ser225 phosphorylation. Lys29 lipid interactions show a slight increase.

Defining the molecular inhibitory mechanism of potent inhibitors like PF-543 is crucial for developing more effective therapeutic modulators. A key finding of this study is the identification of a previously uncharacterized catalytic complex. Probed through three distance pairs—T193–T136, T193–M298, and T136–M298—this complex features a highly dynamic catalytic core involving LBL-1, which wraps around and positions the substrate within the CTD for phosphoryl transfer, alongside the domain interface encompassing the Mg^2+^ATP binding site. This dynamic core appears to transition through intermediates essential for both phosphoryl transfer and release of synthesized S1P. Notably, PF-543—regardless of Mg^2+^ATP presence—shifts the equilibrium among these intermediates, effectively inhibiting catalysis or ligand exchange (Figs. 1 and 6). This inhibitory effect extends to the regulatory loop (Fig. 5 and Supplementary Fig. 6). MD simulations show that PF-543 stabilizes monomeric, membrane-bound SK1 to a degree comparable with Mg^2+^ATP/sphingosine (Supplementary Figs. 7b and 11d). Considering the disengaged R-loop in the crystal template, salt-bridge interactions with the core protein remain intact, resembling those in the unphosphorylated apo and Mg^2+^ATP/sphingosine-bound states (Supplementary Figs. 8b, 8c, 11f, 11g).

Leveraging our ability to distinguish potent from weak SK1 inhibitors, we tested the selective inhibitor SLP7111228 (*K*i = 48 nM), suitable for in vivo studies^26^. Interestingly, like PF-543, SLP7111228 promotes SK1 dimerization, stabilizing a similar conformation but with a slightly lower dimer population (Supplementary Figs. 12a–12c), underscoring dimerization as a shared inhibitory mechanism of potent inhibitors. Unlike PF-543, however, SLP7111228 favors more open lipid-binding loop conformations and, in the presence of Mg^2+^ATP, prevents the fully closed state otherwise stabilized by Mg²⁺ATP/sphingosine (Supplementary Figs. 12d–12e). On the regulatory loop and C-terminal tail, its effect parallels PF-543 (Supplementary Figs. 12f–12k).

PF-543 stabilizes a dimeric configuration similar to that observed in the PF-543 crystal structure (Figs. 1b and 9a), suggesting a functionally relevant state rather than a crystallographic artifact. In this catalytically inactive SK1 state, key membrane-interfacing elements (Leu194, Phe197, and Leu198 in LBL-1) are sequestered within the dimeric interface (Fig. 9a). The presence of a CTD-mediated dimeric structure provides a mechanistic rationale for the CIB1-dependent membrane translocation of SK1, as CIB1 engages SK1 through the same hydrophobic residues in LBL-1 (Fig. 9b). Upon calcium binding, CIB1 undergoes a structural transition that exposes both a membrane-targeting N-terminal myristoyl group and a groove for partner protein interaction—the latter formed by the unfolding of a C-terminal helix (Fig. 9c). Simultaneously, Ser225 phosphorylation partially replaces Asp235 in the R-loop, reconfiguring salt bridges with positively charged residues on the strand pair connecting the two domains (Fig. 9 inset, States 1 to 2; Supplementary Fig. 8). This reconfiguration may induce conformational changes in the strand pair that stabilize a catalytically competent state for sphingosine extraction from the membrane and Mg^2+^ATP binding at the catalytic core (Fig. 9d). Notably, our MD simulations indicate that with phosphorylation and Mg^2+^ATP/sphingosine binding, one of the connecting strands becomes significantly more stable, potentially stabilizing the catalytic complex (Fig. 8b). This strand connects to the SK1 C-terminal tail. Consistently, in our DEER constructs where Ser225 is predominantly phosphorylated or mutated to the phosphomimetic S225D, this R-loop conformation appears particularly stabilized in the presence of MgATP or MgATP/Sph (Figs. 3j and 3l and 3n–3q). Additionally, MD analyses reveal distinct membrane interaction patterns for key membrane-interfacing determinants in phosphorylated versus non-phosphorylated SK1 (Figs. 8c and 8d). Electrostatic interactions between basic residues and anionic lipids are significantly diminished in the MgATP/Sph-bound phosphorylated protein, while hydrophobic interactions increase. In the apo state, Arg186 and Lys27 preferentially interact with SAPI over POPS, with Arg186 also contacting POPE. In the phosphorylated apo state, the hydrophobic patch shows increased contact with POPS. Overall, the lipid contact frequency of the hydrophobic patch and Arg186 remains high across different states, whereas lipid interactions involving Lys27 are significantly reduced, and those involving Lys29 are slightly increased upon Ser225 phosphorylation (Fig. 8d and Supplementary Figs. 11b and 11c). The residues involved in this long-range allosteric transmission will be further investigated through mutational analysis and conformational dynamics studies using DEER spectroscopy. Conserved acidic residues in the C-terminal tail (e.g., Glu381) have also been proposed to interact with the same positively charged residues in the connecting strand pair (Fig. 9 inset, State 3)^14^, thereby stabilizing an inactive conformation—potentially by misaligning key membrane-interacting elements. In this context, Ser225 phosphorylation and R-loop reconfiguration may release the C-terminal tail, a conformation we observe experimentally (Fig. 7), though this observation warrants further investigation. Additionally, the membrane itself may stabilize the NTD-mediated dimeric conformation (Fig. 9e and Supplementary Fig. 2d).

**Figure 9.**
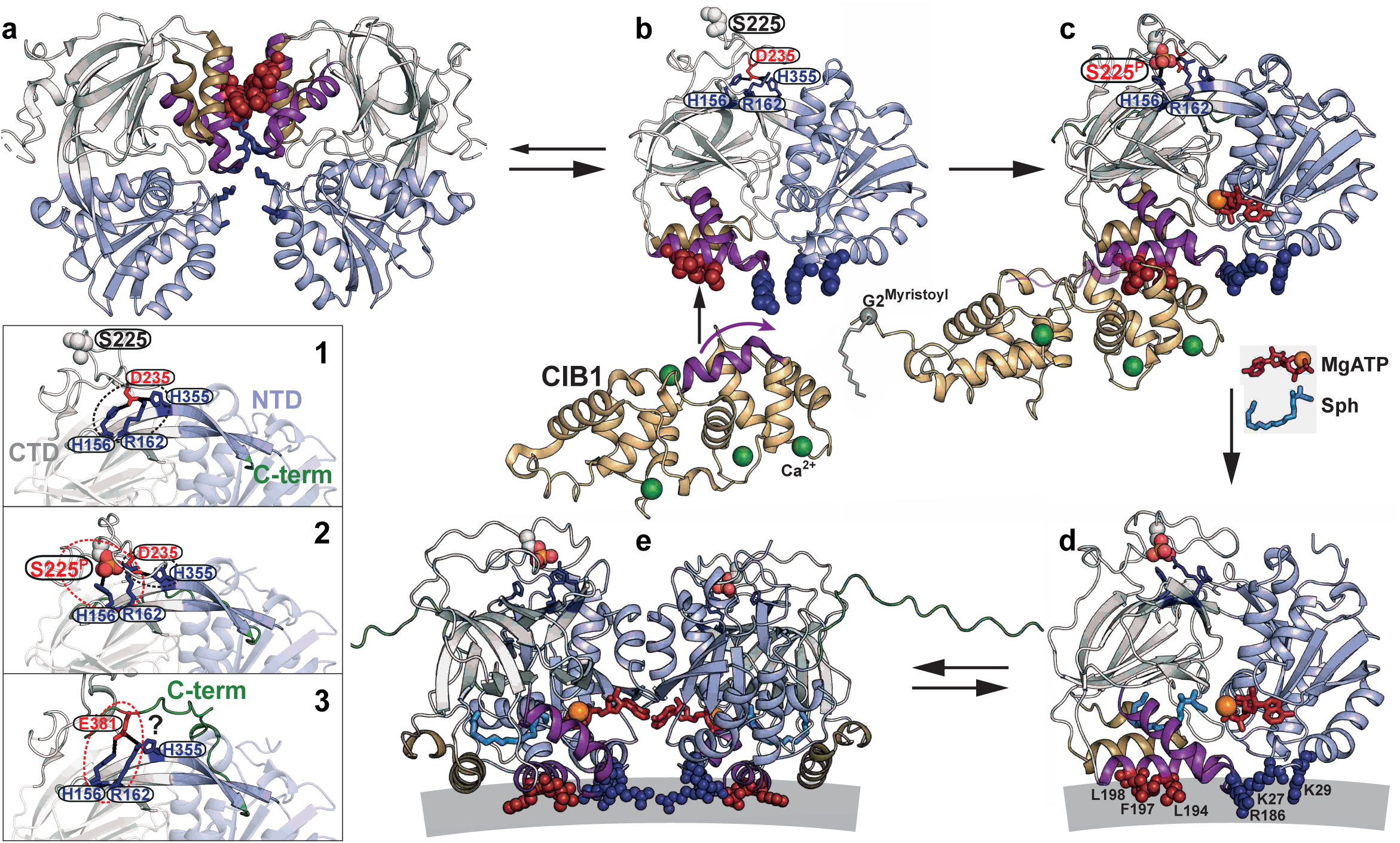
Proposed regulatory mechanism of SK1. (**a**) In its resting state, SK1 may form a CTD-mediated dimer resembling the PF-543–bound crystal structure (PDB code 4V24), representing a catalytically inactive conformation in which key membrane-interacting residues (Leu194, Phe197, and Leu198 in LBL-1; red spheres) are buried within the dimer interface. (**b**) CIB1 (PDB code 1XO5) binds monomeric SK1 (PDB code 3VZB-A) by recognizing these now-exposed hydrophobic residues in LBL-1. Upon calcium binding, CIB1 undergoes a structural transition that exposes its N-terminal myristoyl group for membrane targeting and generates a groove for SK1 interaction through unfolding of a C-terminal helix (**c**; AF3 model). Simultaneously, phosphorylation of Ser225 partially displaces Asp235 in the R-loop, reshaping the salt-bridge network with positively charged residues in the strand pair linking the NTD and CTD (inset: transition from State **1** [PDB code 3VZB-A] to State **2** [AF3 model]). This rearrangement promotes conformational changes that stabilize a catalytically active state for sphingosine extraction and Mg^2+^ATP binding (**d**; AF3 model). Conserved acidic residues in the C-terminal tail (e.g., Glu381) may also engage the same positively charged residues in the strand pair (inset: State **3**; AF3 model), stabilizing an inactive conformation—potentially by misaligning membrane-interfacing elements. In this context, Ser225 phosphorylation and R-loop rearrangement may facilitate release of the C-terminal tail, a conformation supported by our experimental data. (**e**) The membrane itself may further stabilize an NTD-mediated dimeric conformation (AF3 model superimposed on PDB code 3VZC-B/C).

Together, our findings present a unified mechanistic framework for SK1 regulation and inhibition, revealing multilayered regulatory mechanisms governed by conformational flexibility and allosteric transitions. This study not only resolves some of the long-standing questions about SK1 function but also highlights novel therapeutic strategies, including orthosteric inhibition via conformational trapping by driving SK1 into an inactive dimeric state. By distinguishing potent from weak SK1 inhibitors, this study paves the way to screen new compounds, pinpoint strong candidates, and uncover their inhibitory mechanisms with major therapeutic potential.

Integrated with existing structural and cell biology studies, our results offer a comprehensive view of SK1’s regulatory landscape—insights that may also extend to its isoform, SK2, another clinically relevant target.

## Methods

No statistical methods were used to predetermine sample size. The experiments were not randomized. The investigators were not blinded to allocation during experiments and outcome assessment.

### Site-directed mutagenesis

Codon-optimized human SK1 (GenScript) was cloned into a pFastBac-1 vector encoding an N-terminal 6×His tag (44.9 kDa). The seven cysteine residues in SK1 were mutated to alanine via site-directed mutagenesis using complementary oligonucleotide primers, yielding the CL variant. This construct served as the template for introducing single cysteine or double-cysteine pairs and background mutations. Substitution mutations were generated via single-step PCR, in which the entire plasmid was replicated from a single mutagenic primer. SK1 mutants were verified by sequencing with both pFastBac forward and reverse primers (Azenta) to confirm the desired mutations and absence of off-target changes. Mutants are designated by the native residue and sequence position, followed by the substituted residue.

### Expression, purification, and labeling of SK1

Wild-type (WT), cysteine-less (CL), and single-and double-cysteine SK1 mutants were expressed using standard Invitrogen protocols with the pFastBac system. *Spodoptera frugiperda* Sf9 cells (Gibco, ThermoFisher Scientific) were infected at a density of 2–2.5 × 10^6^ cells/mL and incubated for 66–72 hours at 27 °C in a shaking incubator using 2 L disposable flasks. Cells were harvested by centrifugation and stored at −80 °C. For lysis, cell pellets were resuspended in buffer (50 mM Tris⋅HCl, pH 7.8, 200 mM NaCl, 20 mM imidazole, and 10% [vol/vol] glycerol) at 2.6 mL per gram of cells. The buffer was supplemented with 0.5 mM TCEP, 0.9 mM (0.1% [vol/vol]) Tween 20 (Sigma), one EDTA-free protease inhibitor cocktail tablet (Roche), and 1 mM PMSF. The suspension was lysed by sonication, and cell debris was removed by centrifugation at 96,000 × g for 1 hour. The supernatant was incubated with 1.5 mL (bed volume) of His60 Ni-IDA Superflow resin (Takara) at 4 °C for 1.5 hours. After washing with 10 bed volumes of lysis buffer containing 0.9 mM Tween 20, SK1 was eluted using buffer with 250 mM imidazole and 0.9 mM Tween 20. Cysteine mutants were labeled with two rounds of a 10-fold molar excess of 1-oxyl-2,2,5,5-tetramethylpyrroline-3-methyl methanethiosulfonate (Enzo Life Sciences) per cysteine on ice in the dark over a 2-hour period. The samples were then incubated on ice at 4 °C overnight (∼15 hours) to yield the spin-labeled side chain R1. Unreacted spin label was removed by size-exclusion chromatography using a Superdex 200 Increase 10/300 GL column (GE Healthcare) equilibrated in 50 mM Tris⋅HCl, pH 7.8, 200 mM NaCl, and 10% (vol/vol) glycerol, with or without 0.9 mM Tween 20. Peak fractions of purified SK1 were pooled and concentrated using an Amicon Ultra 10,000 MWCO filter concentrator (Millipore), and the final protein concentration was determined by A280 measurement (ε = 49,390 M^-1^⋅cm^-1^) for use in subsequent studies.

### Fluorescence-based sphingosine kinase assay

The functional integrity of spin-labeled DEER mutants was evaluated using a fluorescence-based sphingosine kinase assay, as previously described, with 15-NBD-sphingosine serving as the substrate^40^. Reactions were carried out in a buffer containing 50 mM HEPES (pH 7.4), 15 mM MgCl_2_, 10 mM KCl, 0.005% Triton X-100, and 10% glycerol. The reaction mixture consisted of 20 μM NBD-sphingosine and 1 mM ATP in a final volume of 100 μL. Reactions were initiated by the addition of 1 μM of each SK1 mutant and incubated at 37 °C for 30 minutes. Reactions were quenched by the addition of 100 μL of 1 M potassium phosphate buffer (pH 8.5), followed by extraction with 500 μL of chloroform/methanol (2:1). After centrifugation at 15,000 rpm for 1 minute, 100 μL of the upper aqueous layer was transferred to a 96-well microplate and mixed with 100 μL of dimethylformamide. Fluorescence was measured on a SpectraMax i3 microplate reader with excitation at 485 nm and emission at 538 nm. A reaction lacking enzyme served as a blank control. Each assay was performed in triplicate (technical replicates). For each mutant, the amount of generated NBD-S1P was calculated and normalized to the value obtained for cysteine-less SK1.

### ATPase activity assay

ATPase activity was measured using the malachite green method, as previously described, with Biomol^®^ Green reagent used to detect the release of free inorganic phosphate (P_i_)^41^. Reactions were carried out in a buffer containing 50 mM MOPS (pH 7.4), 10 mM NaCl, and 10 mM MgCl_2_. ATP (100 μM) was added to 50 μL of reaction mixture and transferred to a 96-well plate. Reactions were initiated by adding an equal volume of each SK1 mutant (prepared in the same buffer), resulting in a final SK1 concentration of 0.5 μM per well. Plates were incubated at room temperature for 1 hour. Each SK1 mutant was tested in triplicate. Control reactions lacking ATP were included to account for background signal. Reactions were terminated by adding 200 μL of Biomol^®^ Green reagent per well, followed by incubation at room temperature for 30 minutes to allow color development. Absorbance was measured at 620 nm using a SpectraMax i3 microplate reader. For each mutant, the amount of released phosphate was calculated from a standard curve and normalized to the value obtained for cysteine-less SK1.

### SK1-liposome sample preparation

Cholesterol, 1-palmitoyl-2-oleoyl-sn-glycero-3-phosphocholine (POPC), 1-palmitoyl-2-oleoyl-sn-glycero-3-phosphoethanolamine (POPE), 1-palmitoyl-2-oleoyl-sn-glycero-3-phospho-L-serine (POPS), and L-α-phosphatidylinositol (Liver, Bovine, SAPI) (Avanti Polar Lipids) were combined in a 20:14:35:22:9 molar ratio, dissolved in chloroform, evaporated to dryness using a rotary evaporator, and desiccated overnight under vacuum in the dark. The dried lipids were rehydrated in 50 mM Tris⋅HCl (pH 7.5), 100 mM NaCl, and 10% glycerol buffer to a final concentration of 40 mM, followed by homogenization through 10 freeze-thaw cycles. The resulting lipid suspension was aliquoted and stored at −80 °C. Large unilamellar vesicles (LUVs) were prepared by sequential extrusion of the homogenized lipids through polycarbonate membranes (Avanti) with pore sizes of 0.4, 0.2, and 0.1 μm, performing ≥10 passes through each membrane. SK1 mutants were mixed with liposomes at a 1:1500 molar ratio and concentrated using a 10,000 MWCO filter concentrator to increase total spin concentration.

### CW-EPR and DEER spectroscopy

CW-EPR spectra of spin-labeled SK1 samples were collected at room temperature on a Bruker EMX spectrometer operating at X-band frequency (9.5 GHz) using 10-mW incident power and a modulation amplitude of 1.6 G. Most DEER measurements were performed on an Elexsys E580 EPR spectrometer (Bruker BioSpin) operating at Q-band (33.9 GHz) using the dead-time-free four-pulse sequence at 83 K^48^. Probe pulse lengths were 20 ns (π/2) and 40 ns (π), with a 40-ns pump pulse and a pump–probe frequency separation of 63 MHz. For five DEER pairs (T99–A224, T136– V366, T136–W372, V366, and W372), measurements were performed on a Fathom EPR spectrometer (High Q Technologiesoperating at X-band (9.6 GHz) using the DEER-Stitch method, which combines pulse sequences with 16-step phase cycling, at 50 K. Gaussian probe pulse lengths were ∼15 ns (π/2) and ∼30 ns (π), with a 200-ns adiabatic pump pulse and a pump–probe frequency separation of 80 MHz. Ligands were added in excess relative to protein, yielding final concentrations of 10 mM ATP, 10 mM MgSO4, and 0.55 mM sphingosine, FTY720, SKI-II, PF-543, or SLP7111228. Sample pH was adjusted to 7.4 and verified using a pH microelectrode. For FATHOM and Elexsys DEER measurements, respectively, samples were cryoprotected with 30% and 24% (vol/vol) glycerol and flash-frozen in liquid nitrogen using SQBA sample cartridges and quartz tubes with a 2.1-mm inner diameter and 2.6-mm outer diameter (VitroCom).

All primary DEER decays were analyzed using two complementary approaches: a non-parametric, model-free neural-network method (DEERNet)^49,50^ implemented in DEERanalysis 2026^51^, with each decay analyzed individually, and model-based global Gaussian fitting using a custom MATLAB (MathWorks) program (DeerA; Dr. Richard Stein, Vanderbilt University), as previously described^52^. In the latter approach, DEER decays acquired under different conditions for the same spin-labeled pair were analyzed globally, with the distance distribution represented as a sum of Gaussian components whose number and relative populations were selected according to a statistical criterion. The two approaches yielded comparable distance distributions, and confidence bands were derived from the calculated uncertainties of the fitted parameters. Experimental distance distributions were compared with crystal structures using the rotamer-library approach implemented in MMM 2018.2^53^, with rotamer calculations performed at 175 K.

### Molecular Dynamics simulations

Five systems were prepared for molecular dynamics (MD) simulations: (1) Apo, (2) Apo with Ser225 phosphorylated, (3) Mg^2+^ATP/Sph-bound, (4) Mg^2+^ATP/Sph-bound with phosphorylated Ser225, and (5) PF-543-bound SK1. Each system was simulated in triplicate, with each replicate run for 500 ns. For the non-phosphorylated systems, the crystal structures of SK1 (PDB code 3VZB for systems 1 and 3, and PDB code 4V24 for system 5) were used as starting models^15,16^. For the phosphorylated systems, an AlphaFold 3-predicted model was employed^34^. Histidine residues H156 and H355 were protonated using CHARMM-GUI^54^, based on pKa predictions obtained from PROPKA, with the pH set to 7.4 to mimic physiological conditions. All systems were prepared using the CHARMM-GUI Membrane Builder, which positioned the protein at the membrane interface of a heterogeneous lipid bilayer composed of cholesterol, POPC, POPE, POPS, and SAPI lipids at a molar ratio of 20:14:35:22:9. Membrane positioning was optimized using PPM 2.0_55_. Each protein–membrane complex was solvated in explicit TIP3P water and neutralized with Na⁺ and Cl⁻ ions. An additional concentration of 0.15 M NaCl was added. The final system contained approximately 218,000 atoms.

Simulations were performed with the CHARMM36m all-atom additive force field^56^ using the NAMD 3 simulation engine^57^. Initial energy minimization was conducted for 10,000 steps using conjugate gradient technique. The systems were then equilibrated for approximately 19 ns prior to production runs using a 6-step restraining regimen which was performed in an NVT ensemble based on the CHARMM-GUI procedures for protein simulations^58^. Production MD simulations were run in an NPT ensemble with a 2-fs time step at 310 K using a Langevin integrator with a damping coefficient of 1.0 ps^-1^. Pressure was maintained at 1 atm using the Nose-Hoover Langevin piston method^59,60^. RMSD analyses were carried out with reference to the first frame of the selected trajectory. Salt bridge analysis was based on the minimum distance calculated between donor and acceptor atoms, with a cutoff distance of 4.5 Å used to define a salt bridge. Results were visualized by plotting the minimum distance as a function of simulation time. For lipid contact analysis, a cutoff distance of 4.0 Å was used to define contacts between the protein and lipid molecules.

## Data availability

The generated data, including those from the DEER experiments, are available in the manuscript or the supplementary materials. Source data are provided with this paper. These, including DEER, MD input files, initial and final coordinates, analysis scripts, and analysis data, have been deposited in the Zenodo repository maintained by CERN, https://doi.org/10.5281/zenodo.17059891. Other data that support this study are available from the corresponding author upon request.

## Acknowledgments

The authors thank Mr. Ayush Mistry (Saint Louis University School of Medicine) for assistance with mass photometry and Mr. Nolan McLaughlin and Professor Michael L. Gross (Washington University in St. Louis) for assistance with mass spectrometry. This work was supported by NIH grants R37-CA265877 and S10-OD036230 (to R.D.) and by the Doisy Fund of the Edward A. Doisy Department of Biochemistry and Molecular Biology at the Saint Louis University School of Medicine. The molecular dynamics studies were supported by NIH grant R35-GM147423 (to M.M.). This research used resources of the Oak Ridge Leadership Computing Facility at the Oak Ridge National Laboratory, which is supported by the Office of Science of the U.S. Department of Energy under Contract No. DE-AC05-00OR22725. The authors also acknowledge the Texas Advanced Computing Center (TACC) at The University of Texas at Austin for providing computational resources through LRAC award CHE21003 (to M.M.).

## Author contributions

R.D. and M.M. designed the experiments and simulations. B.A.E. created, expressed, and purified all SK1 mutants and performed the functional integrity analyses. R.D. performed the EPR experiments and analyzed the data. A.S. and H.W. performed and analyzed the simulations. R.D., M.M., and B.A.E. wrote the paper.

## Supplementary Information

**Supplementary Figure 1.**
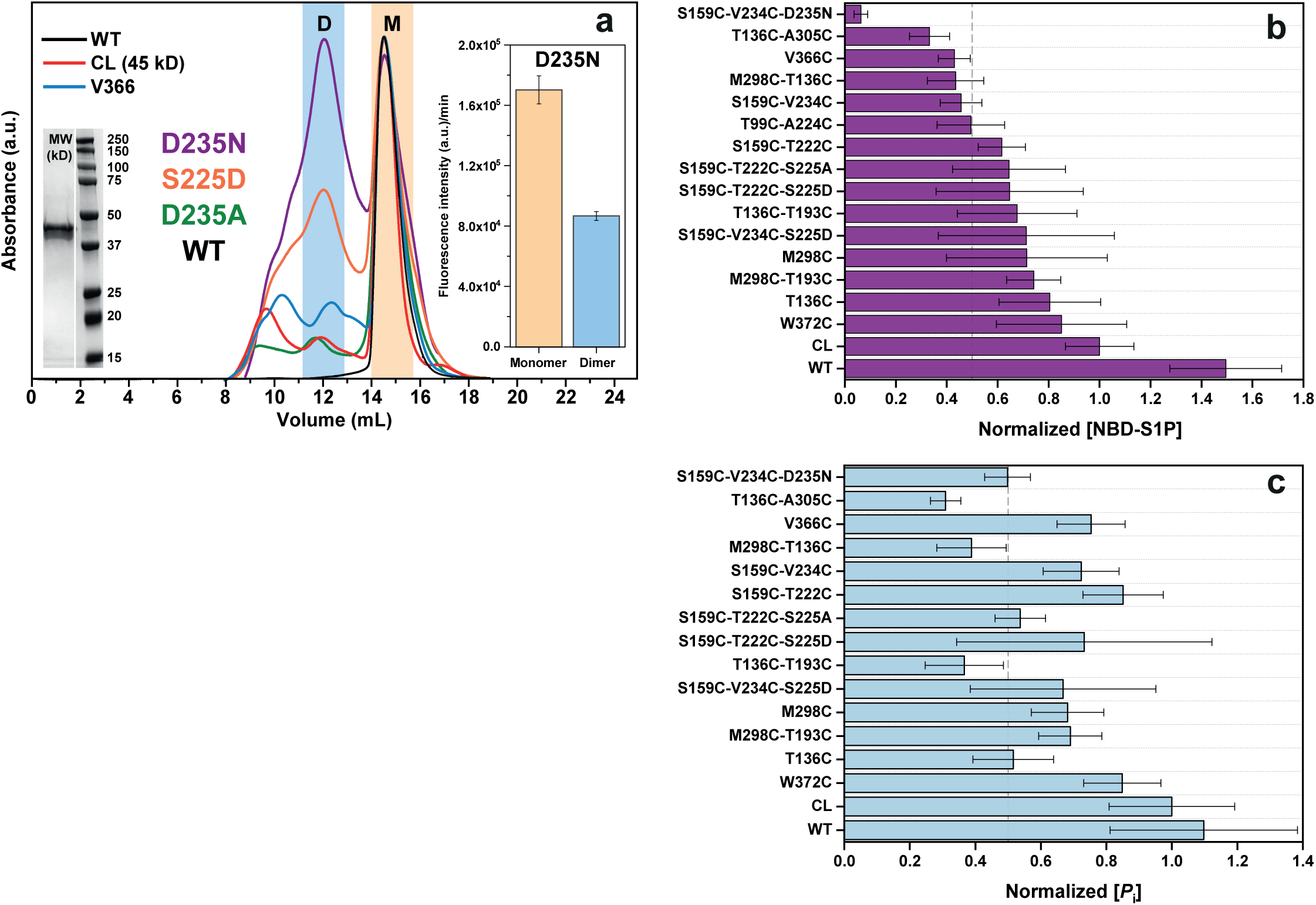
Functional integrity analyses. (**a**) Size-exclusion chromatography profiles of wild-type (WT), cysteine-less (CL), representative spin-labeled SK1 mutants, and the S225D, D235N and D235A variants on the WT background. Dimeric fractions increased in the order D235N > S225D > D235A > WT. Monomeric (beige) and dimeric (blue) fractions of D235N variant were assayed for sphingosine kinase activity using 25 μM NBD-sphingosine (inset). The dimeric fraction exhibited approximately half the activity of the monomeric fraction, supporting a potential regulatory role for SK1 dimerization. **(b)** Fluorescence-based sphingosine kinase activity of WT, CL and spin-labeled DEER mutants, normalized to CL SK1. **(c)** ATPase activity of the same constructs, normalized to CL SK1. Assays were performed in triplicate (technical replicates; *n* = 3). Source data are provided as a Source Data file.

**Supplementary Figure 2.**
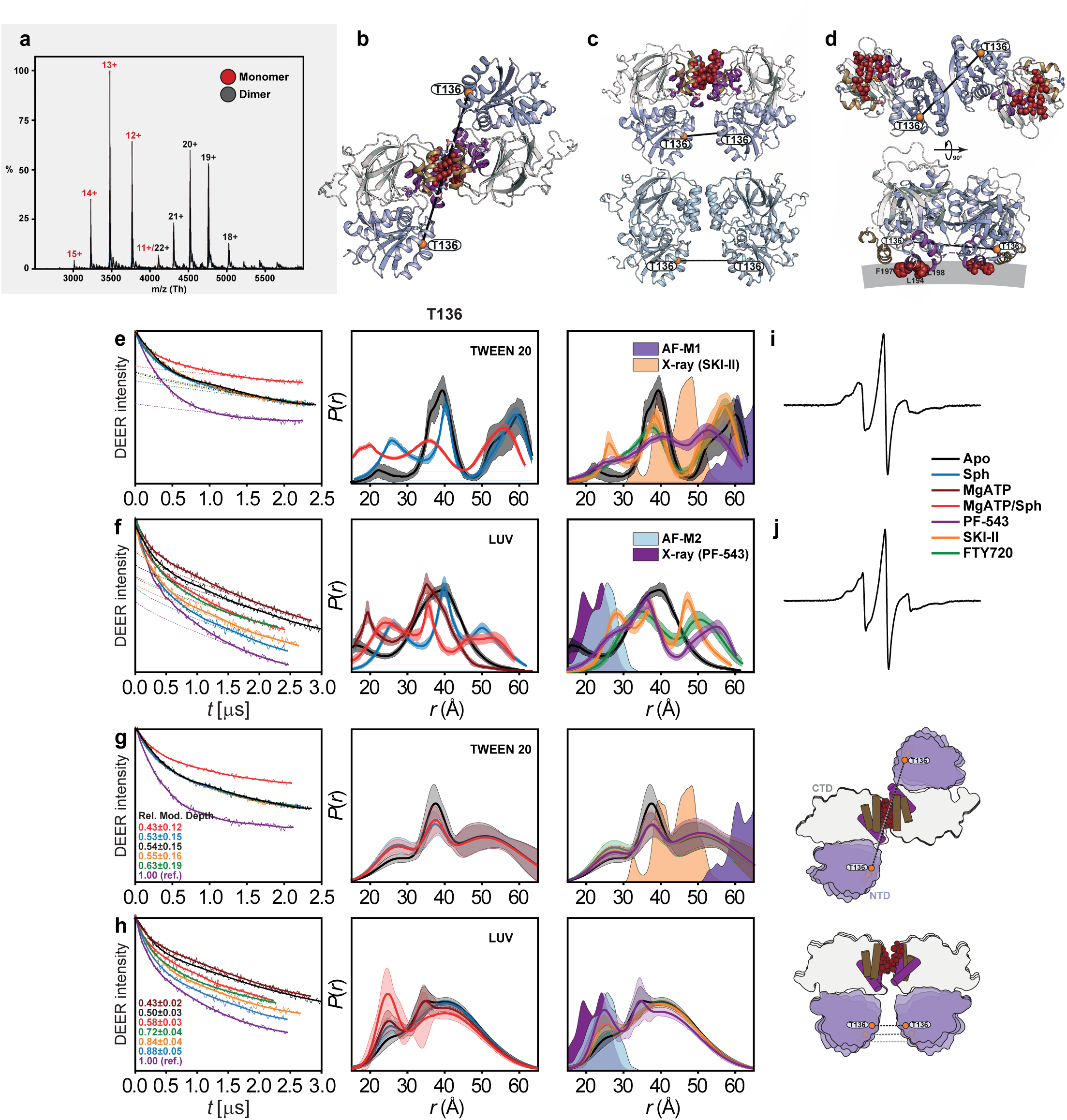
SK1 dimerization is stabilized by CTD/LBL contacts while the NTDs remain dynamic. (**a**) Native mass spectrometry of WT SK1 (20 µM) in detergent-free buffer reveals a monomer–dimer equilibrium, with charge-state envelopes corresponding to ∼45 kDa monomeric species (11+–15+) and ∼90 kDa dimeric species (18+–21+). (**b**–**d**) SK1 dimer models and crystal structures with the NTD, CTD, LBL-1 and LBL-3 colored light blue, white, purple and sand, respectively. Membrane-binding hydrophobic patches are shown as dark-red spheres and spin-labeled T136 sites in the NTD as orange spheres. (**b**) PF-543-bound AF-M1 dimer in Tween 20, featuring rotated protomers stabilized by CTD-mediated LBL contacts. (**c**) PF-543-bound crystal structure (PDB 4V24; top) and parallel AF-M2 dimer stabilized by LBL contacts under detergent-free conditions (bottom). (**d**) SKI-II-bound dimer (PDB 3VZC, chains B/C), illustrating a distinct NTD-mediated interface. (**e**–**h**) DEER measurements at T136 reveal broad inter-NTD distance distributions in both Tween 20 and LUVs, consistent with dynamic NTD orientations and the absence of a stable NTD–NTD interface. Panels **e** and **f** were analyzed using the deep neural network approach DEERNet, whereas panels **g** and **h** were analyzed by global Gaussian fitting. Predicted distance distributions from the corresponding AF and crystal structures are shaded for comparison. The broader distributions observed in LUVs further support enhanced NTD conformational heterogeneity with a membrane environment. PF-543-bound data support CTD/LBL-mediated dimerization, whereas the Mg^2+^ATP/sphingosine catalytic complex shows a lower extent of dimerization (relative modulation depth: 0.43 ± 0.12 in Tween 20 and 0.58 ± 0.03 in LUVs). (**i**,**j**) CW-EPR spectra of T136-labeled SK1 in Tween 20 (**i**) and LUVs (**j**). Together, the structural models and DEER measurements support a dimer architecture stabilized primarily by CTD/LBL contacts, with dynamically oriented NTDs lacking persistent inter-protomer contacts.

**Supplementary Figure 3.**
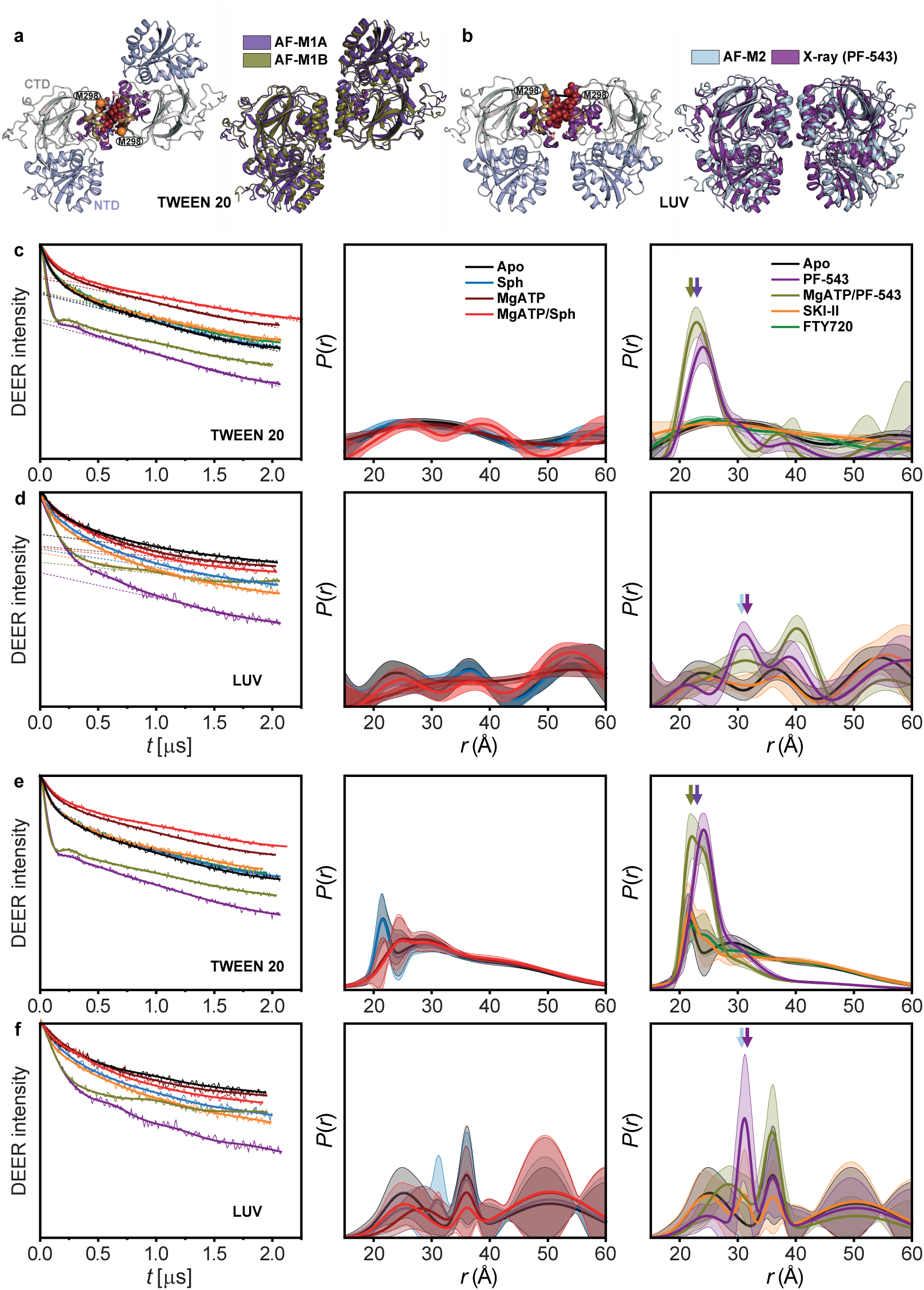
Independent non-parametric and model-based DEER analyses converge on the same SK1 dimerization landscape. Independent analysis of singly labeled SK1 at M298 using non-parametric and global Gaussian-fitting approaches yields closely consistent distance distributions and the same mechanistic interpretation: potent inhibitors stabilize two discrete SK1 dimer architectures mediated by the lipid-binding loops, whereas apo and catalytic conditions, including Mg^2+^ATP/Sph, produce substantially lower levels of dimerization with weaker, more broadly distributed dimeric populations. (**a**) PF-543-bound SK1 dimer model in Tween 20 (AF-M1A, violet), characterized by rotated protomers, aligned with the Mg^2+^ATP/PF-543 model (AF-M1B, olive). (**b**) PF-543-bound crystal structure (PDB 4V24, purple) aligned with the parallel, side-by-side dimer model AF-M2 (light blue). Membrane-binding hydrophobic patches are shown as dark-red spheres and spin-labeled M298 sites on LBL-3 as orange spheres. (**c**,**d**) Primary DEER traces and corresponding distance distributions, *P*(*r*), obtained by DEERLab non-parametric (model-free) analysis^51^ for substrate- and inhibitor-bound conditions in Tween 20 (**c**) and liposomes (LUV; **d**). (**e**,**f**) Corresponding analyses using model-based global Gaussian fitting^52^ in Tween 20 (**e**) and LUVs (**f**). For both approaches, confidence bands indicate uncertainty associated with fitting the primary DEER traces. Both analyses consistently reveal weak, heterogeneous dimerization under apo and catalytic conditions, including Mg^2+^ATP and Mg^2+^ATP/Sph, whereas PF-543 stabilizes well-defined dimeric states. Predicted distance distributions from the PF-543-bound crystal structure, AF-M2, AF-M1/AF-M1A and AF-M1B models are indicated by purple, light-blue, violet and olive arrows, respectively.

**Supplementary Figure 4.**
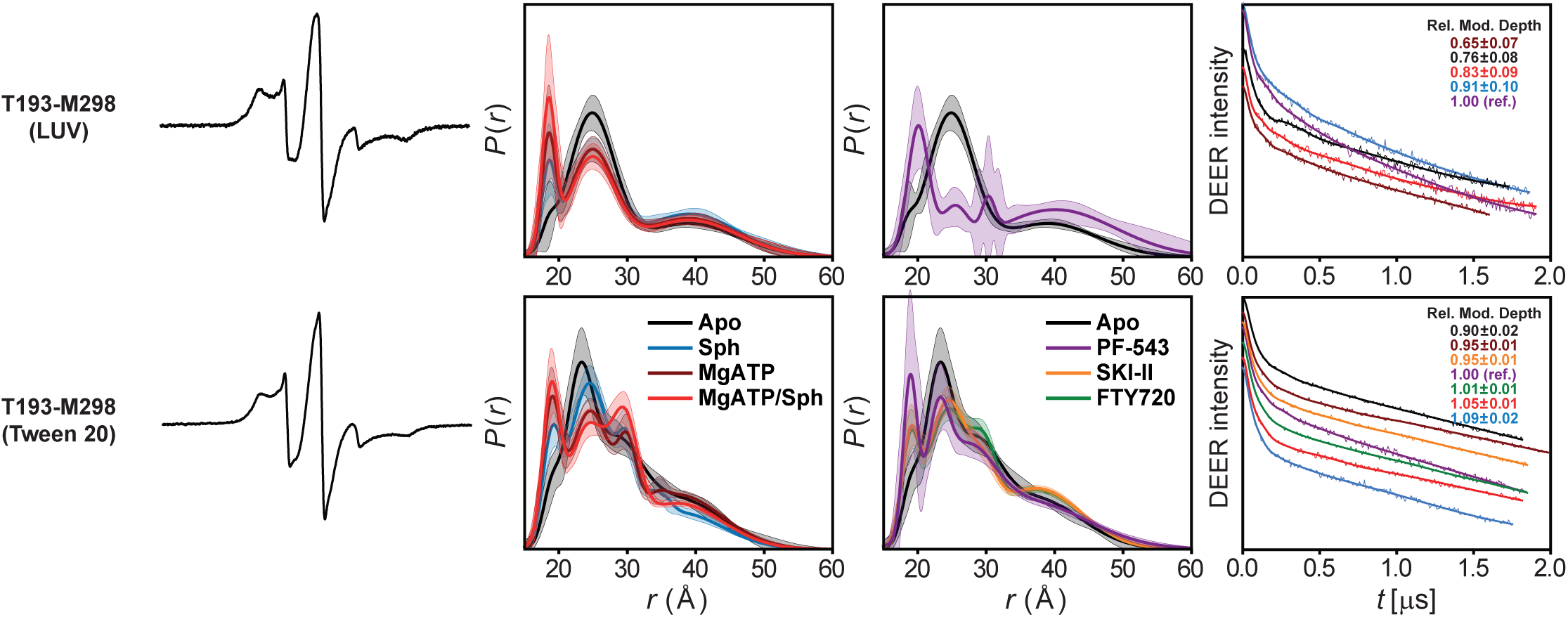
DEER data analysis for distance pairs probing the conformational dynamics of the lipid-binding loops for sphingosine entry and binding. For each mutant, from left to right, CW EPR, global Gaussian-fitted distance distributions with confidence bands (2σ) about the best fit lines, and the primary DEER traces along with the fits are shown.

**Supplementary Figure 5.**
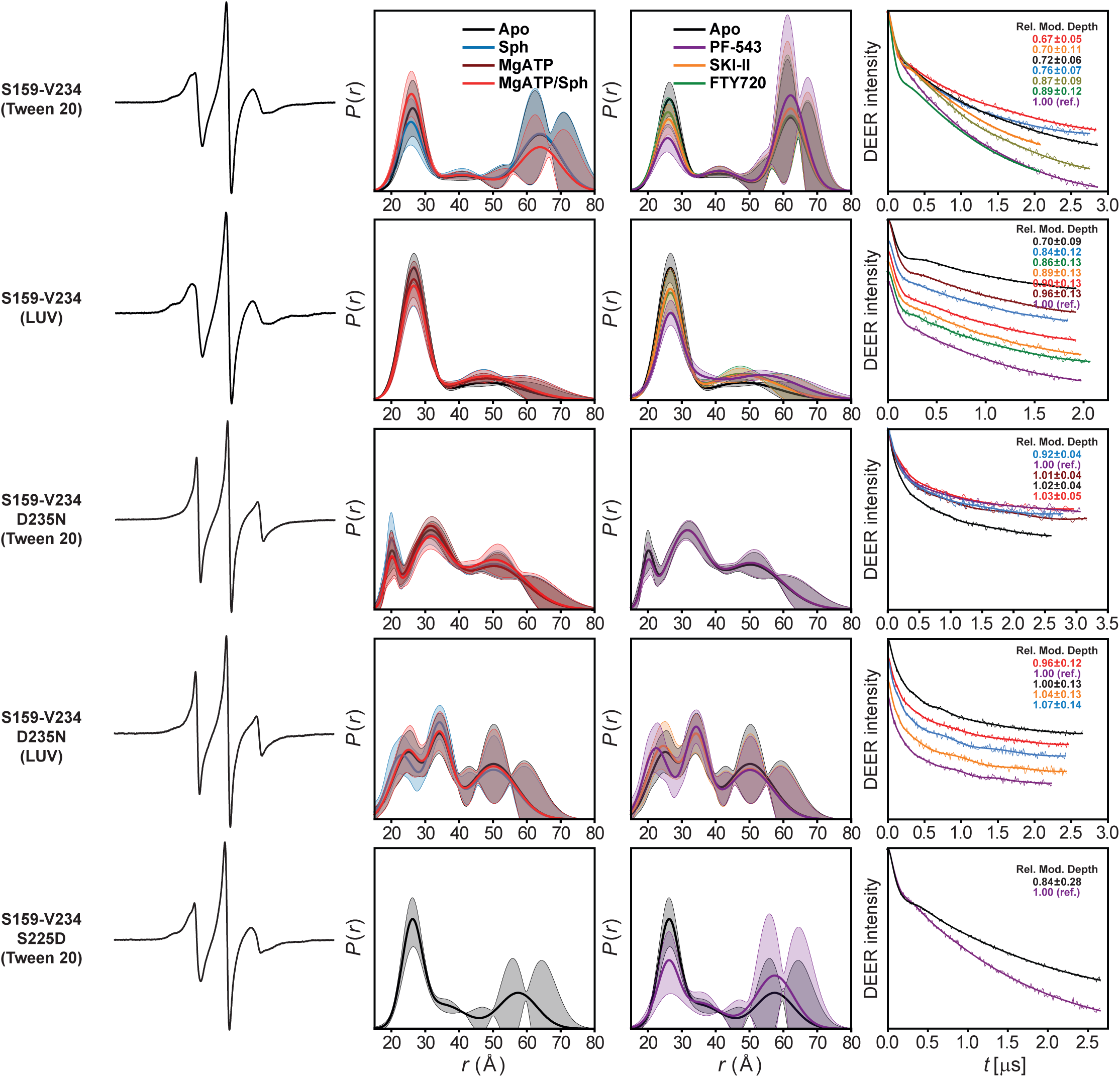
DEER data analysis for distance pair S159-V234, probing the conformational dynamics of the regulatory loop. For each mutant, from left to right, CW EPR, global Gaussian-fitted distance distributions with confidence bands (2σ) about the best fit lines, and the primary DEER traces along with the fits are shown.

**Supplementary Figure 6.**
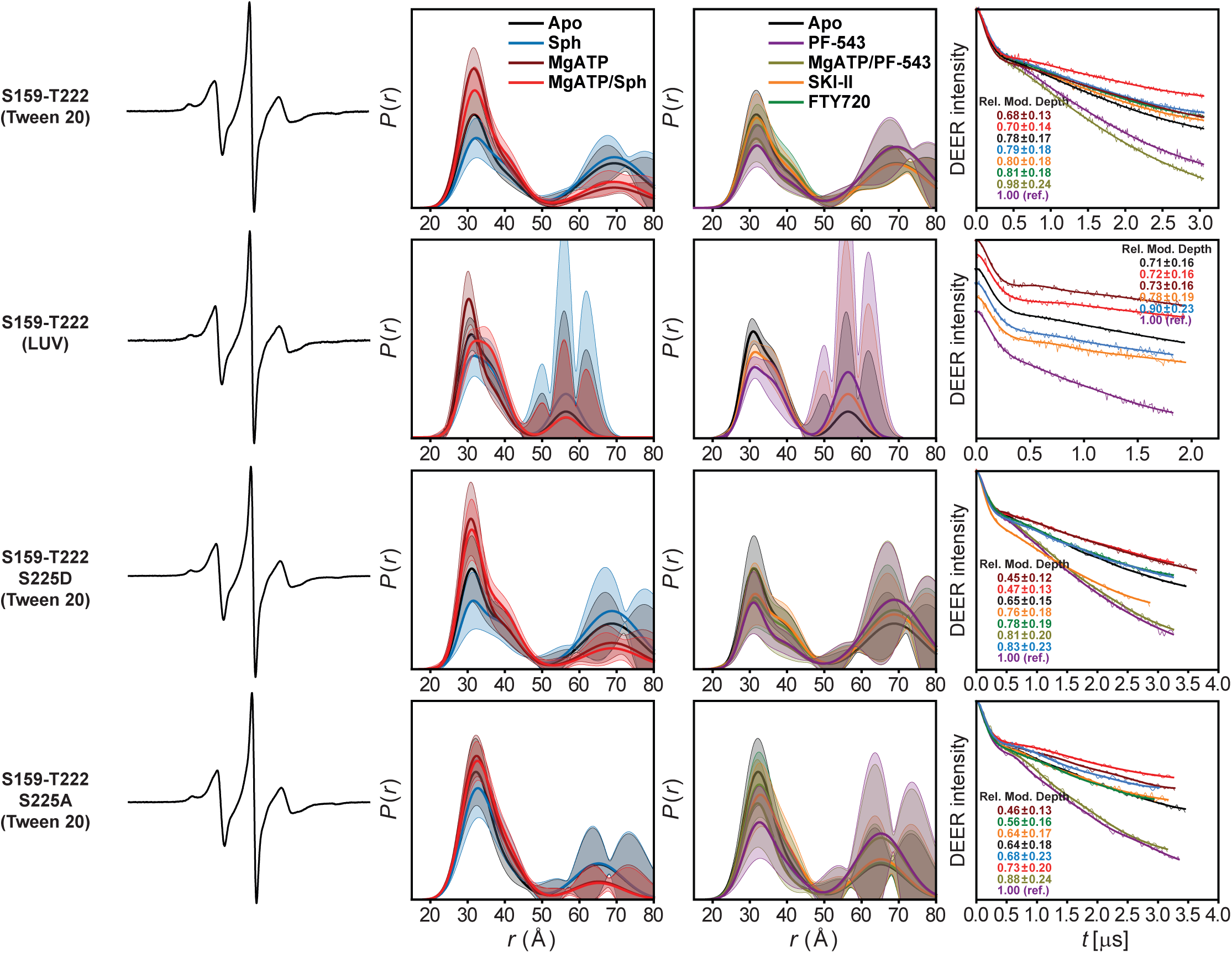
DEER data analysis for distance pair S159-T222, probing the conformational dynamics of the regulatory loop. For each mutant, from left to right, CW EPR, global Gaussian-fitted distance distributions with confidence bands (2σ) about the best fit lines, and the primary DEER traces along with the fits are shown.

**Supplementary Figure 7.**
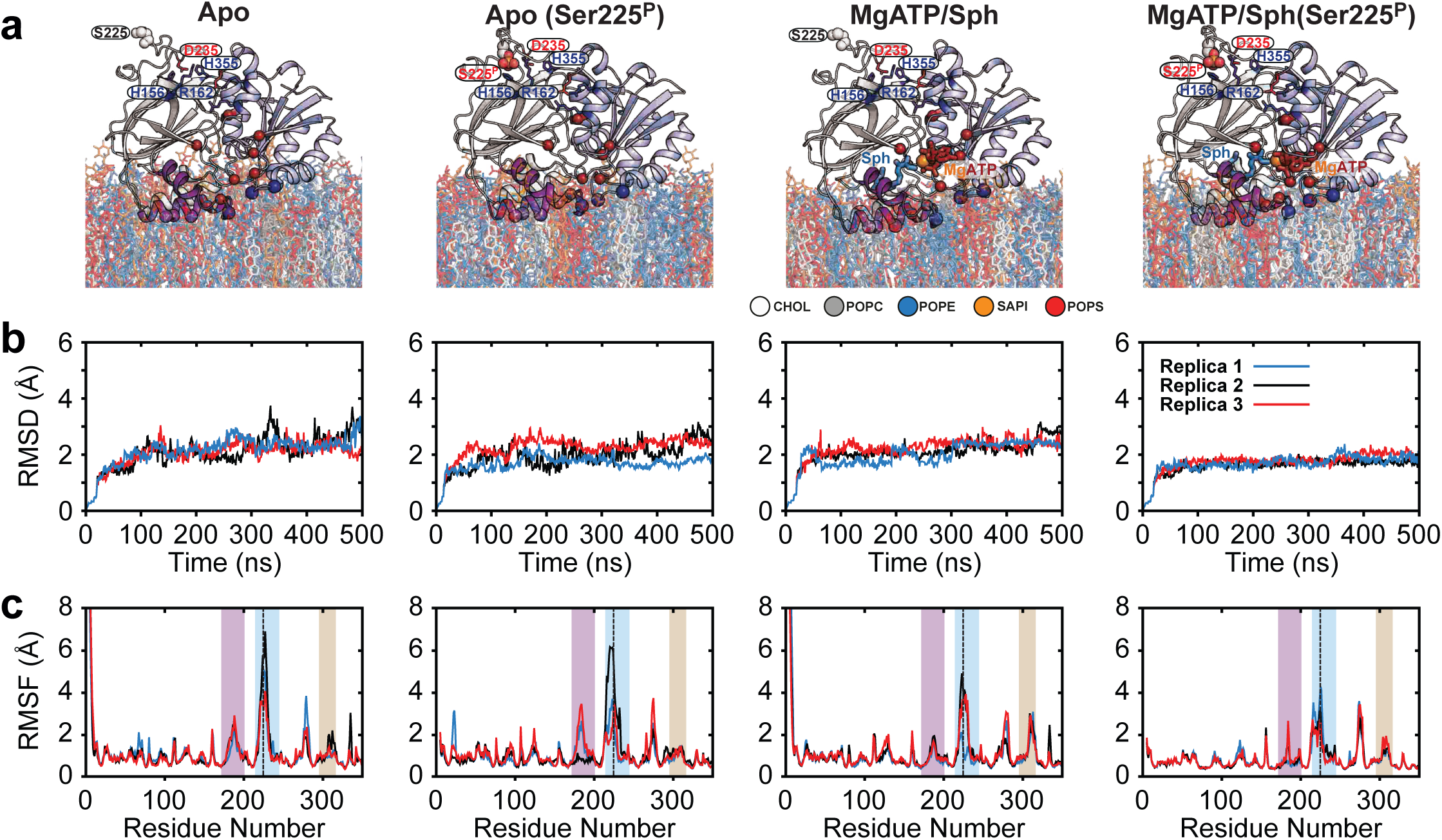
Molecular dynamics simulations of SK1 bound to plasma membrane (PM) lipid bilayers. (**a**) Representative snapshots of four MD simulation systems: Apo, Apo(Ser225^P^), Mg^2+^ATP/Sph, and Mg^2+^ATP/Sph(Ser225^P^). (**b**) Backbone root-mean-square deviation (RMSD) of Cα atoms over 500 ns for three independent replicates of each system, demonstrating overall structural stability. (**c**) Per-residue root-mean-square fluctuation (RMSF) profiles for the same replicates, with shaded purple, blue, and beige regions marking LBL-1, the regulatory loop, and LBL-3, respectively. The phosphorylation site Ser225 is indicated by a dashed line. SK1 remains conformationally stable in the membrane-bound state, with apo systems showing greater flexibility than ligand-bound states. Binding of Mg^2+^ATP/sphingosine, especially with Ser225 phosphorylation, produces the most stable configuration by restraining the regulatory loop. These results reveal that Mg^2+^ATP/sphingosine and Ser225 phosphorylation act together to allosterically stabilize SK1. MD simulations reveal distinct dynamic behaviors of LBL-1 and LBL-3 across states.

**Supplementary Figure 8.**
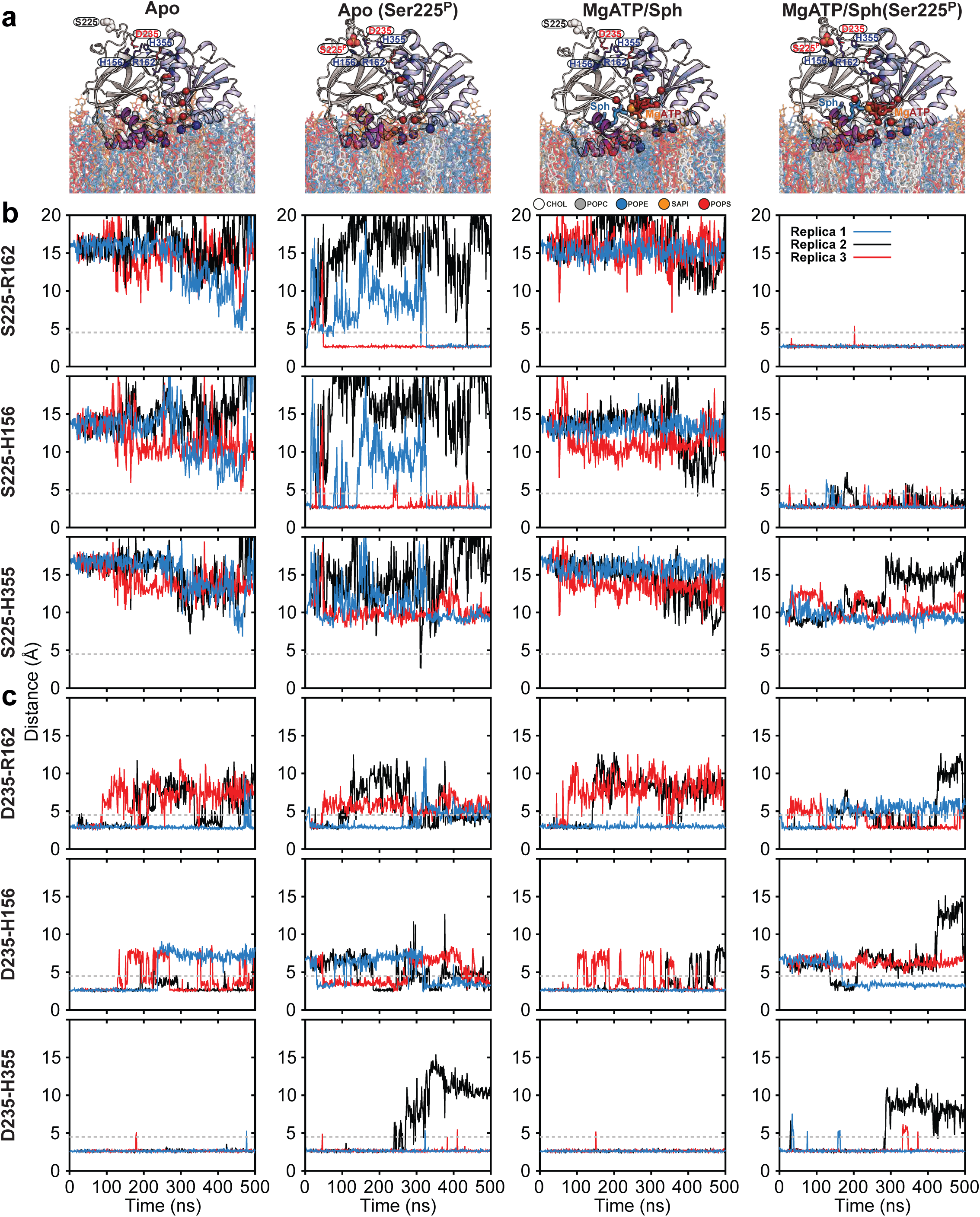
R-loop salt bridges remodeled by phosphorylation and substrates. (**a**) Representative snapshots of four MD simulation systems, each performed in three independent replicates. (**b**,**c**) Time series of the distances between three basic residue side chains and either Ser225/Ser225^P^ or Asp235 during simulations, used to monitor salt bridge formation between the R-loop and the strand pair connecting the NTD to the CTD. Phosphorylation of Ser225 stabilizes salt bridges to His156 and Arg162, reconfiguring the R-loop. Binding of substrates (Mg^2+^ATP and Sph) mutually and allosterically stabilizes this R-loop conformation. Regardless of the condition, Asp235 consistently maintains a tight interaction with His355, while the salt bridge with His156 is less stable and becomes disrupted upon Ser225 phosphorylation.

**Supplementary Figure 9.**
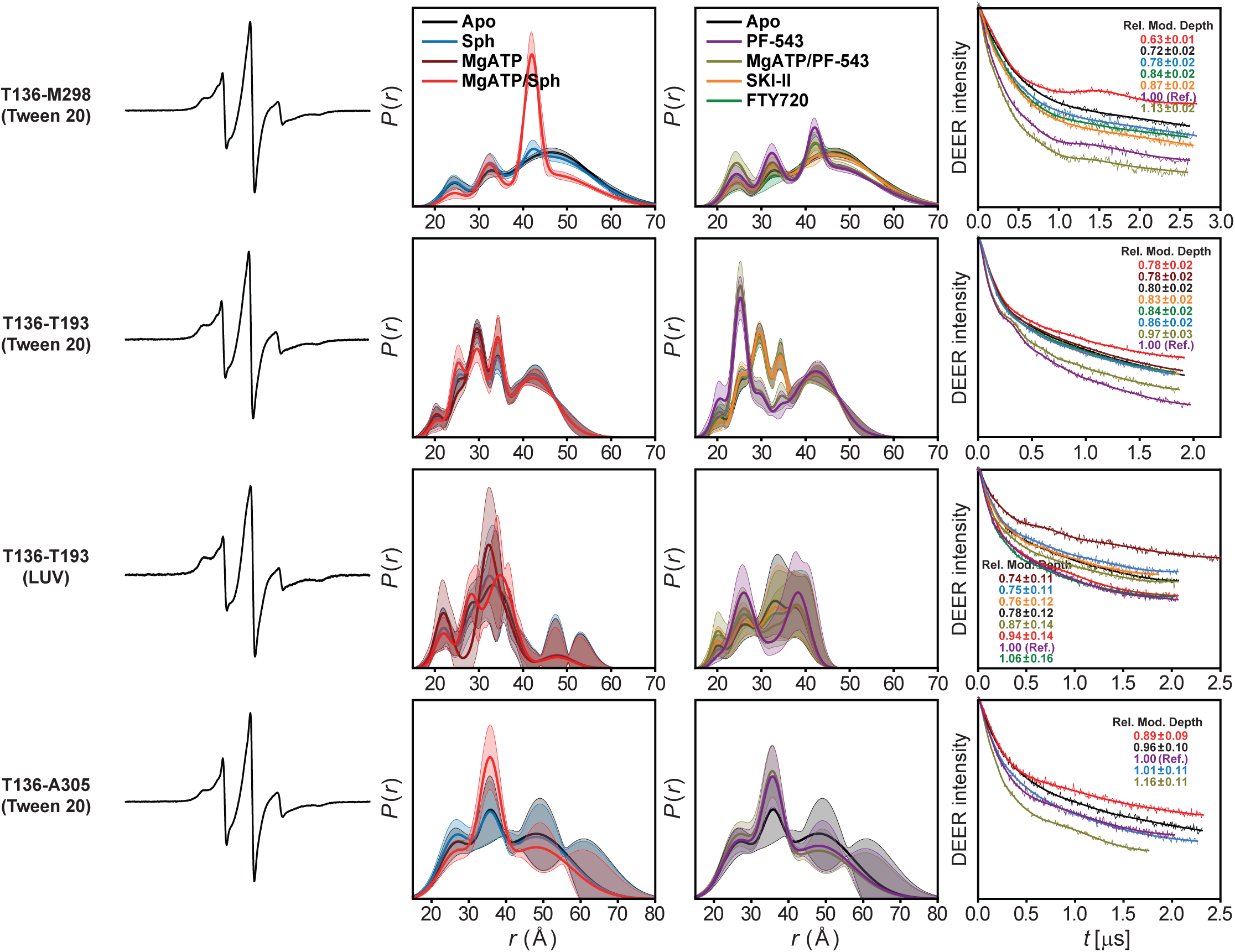
DEER analysis of inter-domain distances reveals NTD–CTD conformational ensembles regulated by substrates, inhibitors, and S1P synthesis. For each mutant, from left to right, CW EPR, global Gaussian-fitted distance distributions with confidence bands (2σ) about the best fit lines, and the primary DEER traces along with the fits are shown.

**Supplementary Figure 10.**
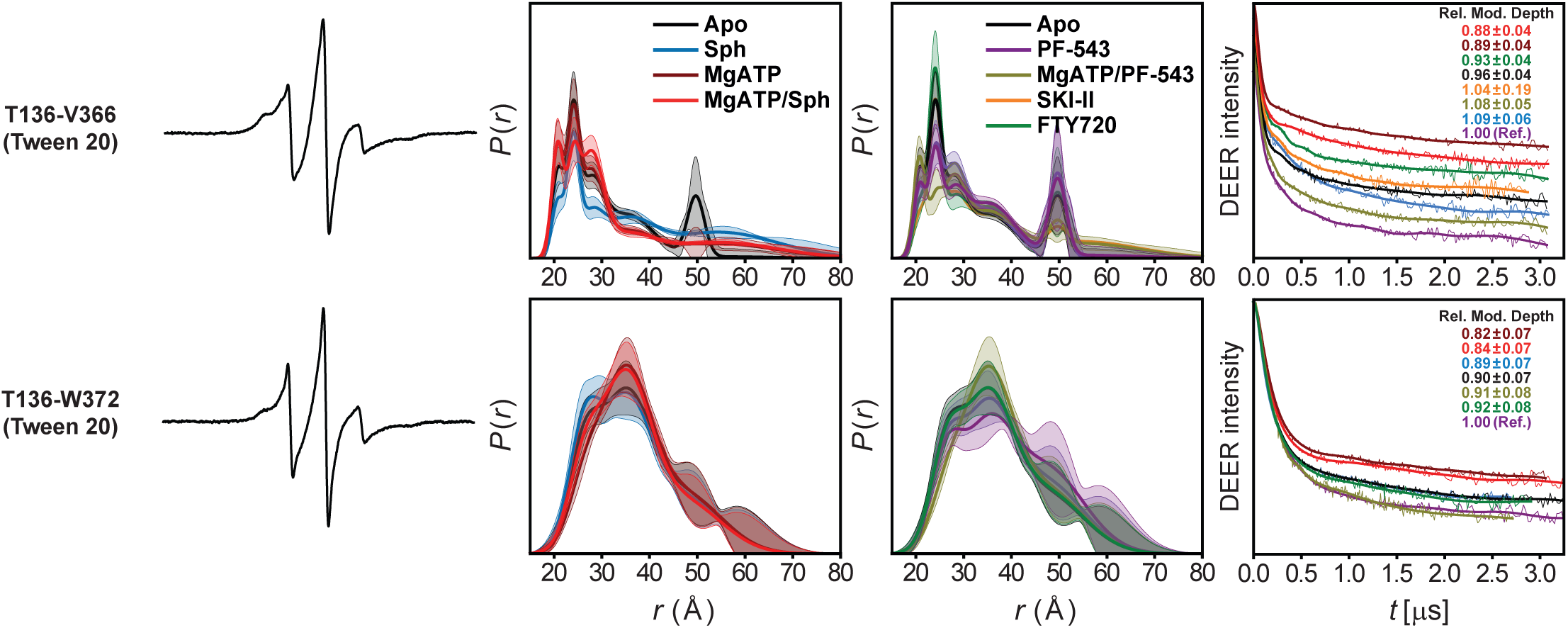
DEER analysis of distance pairs probing the C-terminal tail conformational ensemble highlights differences between proximal and distal segments. For each mutant, from left to right, CW EPR, global Gaussian-fitted distance distributions with confidence bands (2σ) about the best fit lines, and the primary DEER traces along with the fits are shown.

**Supplementary Figure 11.**
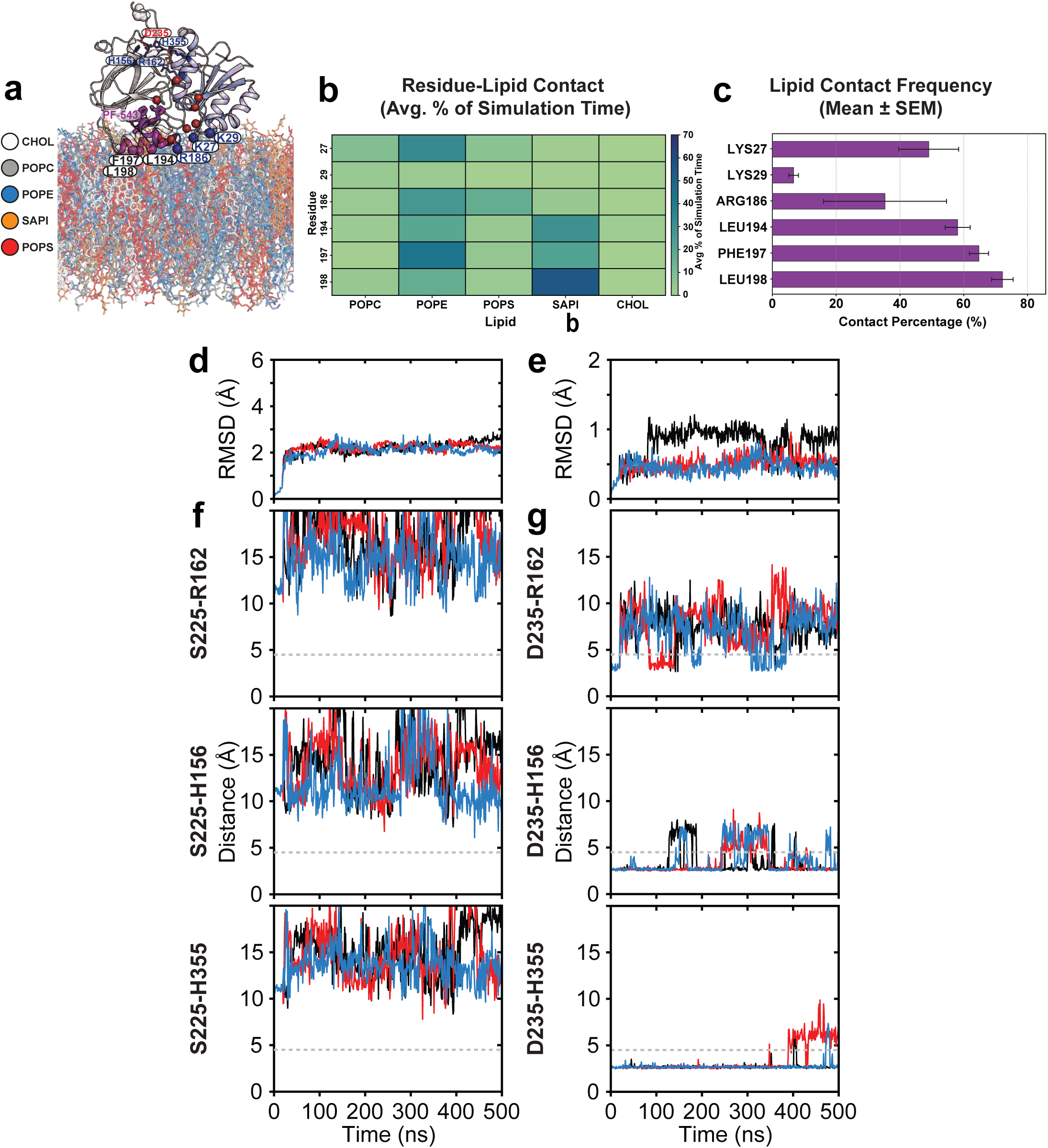
PF-543 mimics Mg^2+^ATP/sphingosine in stabilizing monomeric SK1. (**a**) Representative snapshot of the MD simulation system bound to PF-543, performed in three independent replicates. Membrane-interacting residues are shown as spheres: dark blue (basic) and purple (hydrophobic). (**b**) Lipid interaction specificity of membrane-interfacing residues, plotted as the average contact percentage over the total simulation time, reveals a distinct interaction pattern for the PF-543–bound protein compared with other systems (Fig. 8). Electrostatic interactions between basic residues and anionic lipids are largely diminished, except for moderate Arg186–POPS contacts, while hydrophobic residues show strong engagement with SAPI and POPE. Notably, POPE exhibits the highest lipid contacts across all species. (**c**) Overall, the lipid contact frequency pattern of the PF-543–bound system resembles that of the Mg^2+^ATP/sphingosine system (Fig. 8d), with extensive membrane interactions but slightly reduced Arg186 contacts. (**d**) Backbone root-mean-square deviation (RMSD) of Cα atoms over 500 ns for three independent replicates of each system indicates that PF-543 stabilizes monomeric, membrane-bound SK1 comparably to Mg^2+^ATP/sphingosine. (**e**) Flexibility of the NTD/CTD connecting strand (residues 350–362) in PF-543–bound SK1 is similar to the apo system (Fig. 8b), consistent with the disengaged Ser225 in both systems. (**f**,**g**) Time series of distances between three basic residue side chains and either Ser225 or Asp235 during simulations, monitoring salt-bridge formation between the R-loop and the NTD–CTD connecting strand. Despite the disengaged R-loop observed in the crystal template, core salt-bridge interactions remain intact, resembling those in apo and Mg^2+^ATP/sphingosine-bound states.

**Supplementary Figure 12.**
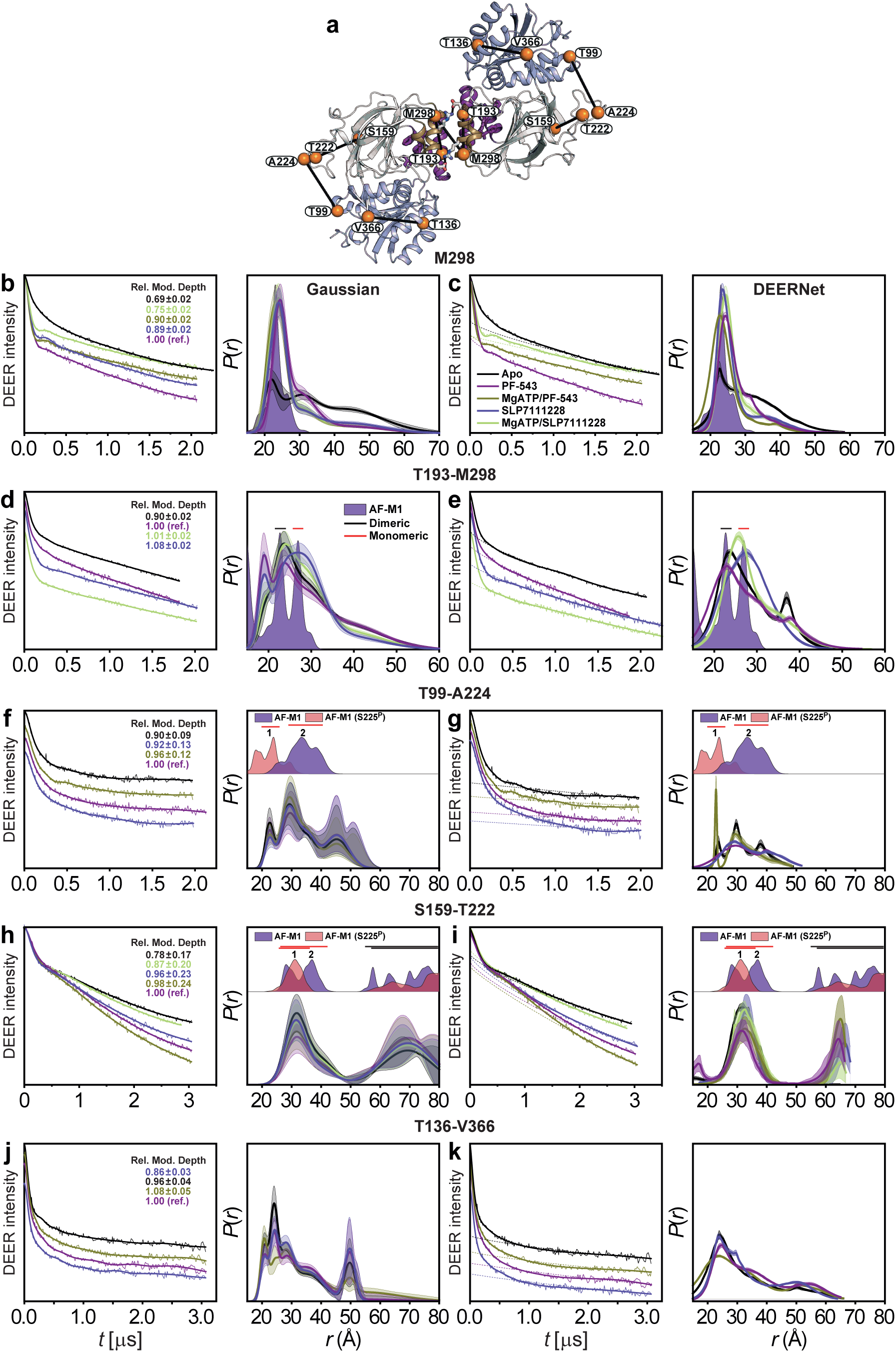
Dimerization as a shared inhibitory mechanism of potent inhibitors. (**a**) Cartoon representation of the PF-543–bound SK1 dimer model in Tween 20 (AF-M1) with rotated monomers, showing the NTD (light blue), CTD (white), LBL-1 (purple), and LBL-3 (sand); spin-labeled sites are highlighted as orange spheres. Primary DEER traces with fits are shown for (**b**,**c**) singly labeled SK1 at M298 to assess dimerization, (**d**,**e**) the T193–M298 pair (LBL-1 to LBL-3) to probe LBL conformational ensembles, (**f**–**i**) the T99–A224 and S159–T222 pairs to monitor changes near Ser225 on the regulatory loop, and (**j**,**k**) the T136–V366 pair probing the C-terminal tail conformational ensemble, in the presence of ligands and inhibitors with Tween 20. Corresponding distance distributions *P*(*r*) with 2σ confidence bands are shown; these bands reflect uncertainty associated with fitting the primary DEER traces. The potent inhibitor SLP7111228 (*K*_i_ = 48 nM) promotes dimerization similar to PF-543 but with a slightly reduced dimer population, underscoring dimerization as a shared inhibitory strategy. Unlike PF-543, SLP7111228 favors more open lipid-binding loop conformations and, in the presence of Mg^2+^ATP, prevents formation of the fully closed state stabilized by Mg^2+^ATP/sphingosine. Its effects on the regulatory loop and C-terminal tail closely mirror those of PF-543, although the regulatory-loop response is modestly attenuated.

**Supplementary Figure 1a.**
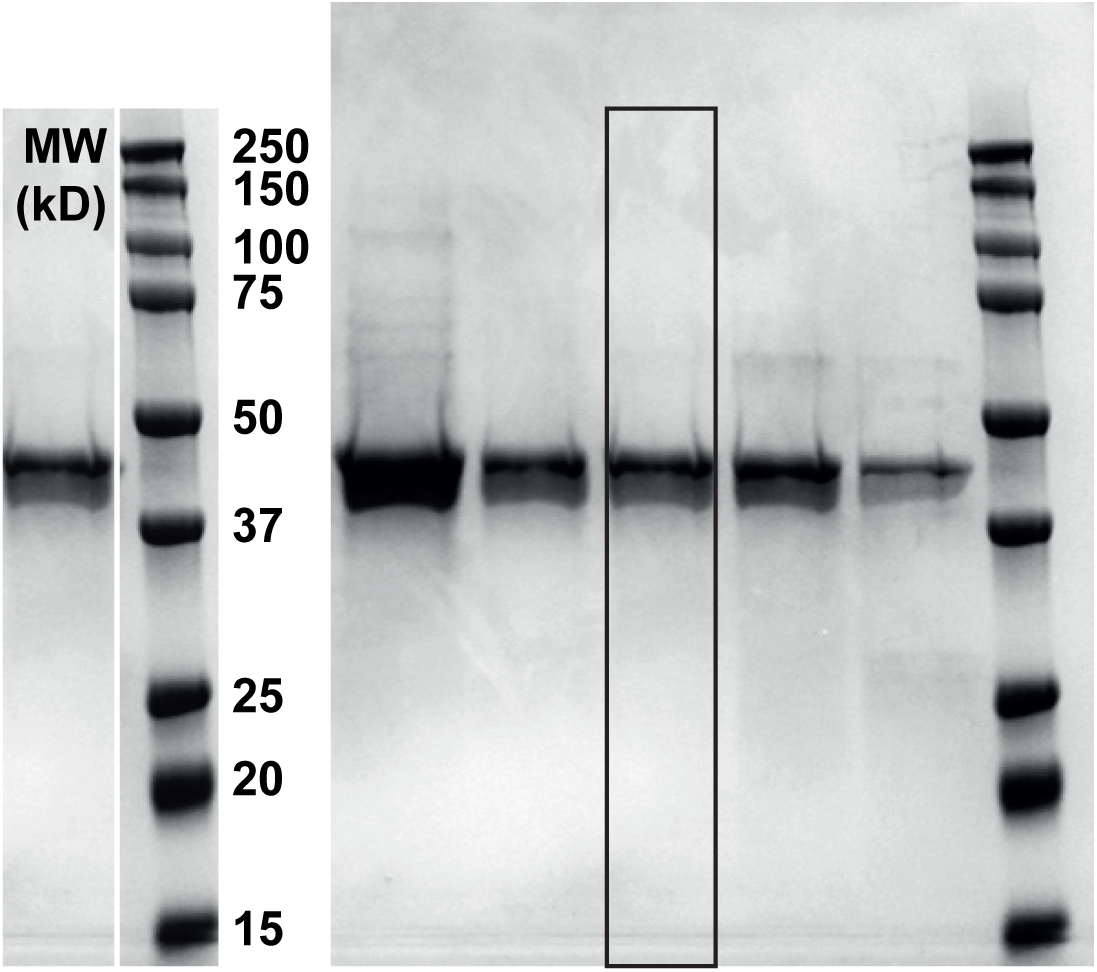

